# Clonal stochasticity in early NK cell response to mouse cytomegalovirus is generated by mature subsets of varying proliferative ability

**DOI:** 10.1101/2023.09.07.556760

**Authors:** Darren Wethington, Saeed Ahmad, Marc Potempa, Giuseppe Giuliani, Oscar A. Aguilar, Maheshwor Poudel, Simon Grassmann, William Stewart, Nicholas M. Adams, Joseph C. Sun, Lewis L. Lanier, Jayajit Das

## Abstract

Natural killer (NK) cells are classically defined as innate immune cells, but experiments show that mouse cytomegalovirus (MCMV) infection in C57BL/6 mice can cause NK cells to undergo antigen-specific proliferation and memory formation, similar to adaptive CD8+ T cells. One shared behavior between CD8+ T cells and NK cells is clonal expansion, where a single stimulated cell proliferates rapidly to form a diverse population of cells. For example, clones derived from single cells are most abundant during expansion when they are primarily CD27- for NK cells and CD62L- for T cells, phenotypes derived from precursor CD27+ and CD62L+ cells, respectively. Here we determined the mechanistic rules involving proliferation, cell death, and differentiation of endogenous and adoptively transferred NK cells in the expansion phase of the response to MCMV infection. We found that the interplay between cell proliferation and cell death of mature CD27- NK cells and a highly proliferative CD27-Ly6C- mature subtype and intrinsic stochastic fluctuations in these processes play key roles in regulating the heterogeneity and population of the NK cell subtypes. Furthermore, we estimate rates for maturation of endogenous NK cells in homeostasis and in MCMV infection and found that only NK cell growth rates, and not differentiation rates, are appreciably increased by MCMV. Taken together, these results quantify the differences between the kinetics of NK cell antigen-specific expansion from that of CD8+ T cells and unique mechanisms that give rise to the observed heterogeneity in NK cell clones generated from single NK cells in the expansion phase.

## Introduction

Natural Killer (NK) cells are lymphocytes of the innate immune system and provide important protection against viral infections and tumors (Caligiuri, 2008; Vivier *et al*, 2008). NK cells develop in the bone marrow and express a diverse set of germline-encoded activating and inhibitory receptors (Abel *et al*, 2018; Biassoni, 2008; Huntington *et al*, 2007). Mouse cytomegalovirus (MCMV) infection induces antigen-specific expansion of naïve Ly49H+ NK cells in mice, followed by a contraction phase, and then the formation of long-lived adaptive NK cells displaying recall responses to the same antigen (Cerwenka & Lanier, 2016; Robbins *et al*, 2004; Schlub *et al*, 2011; Sun *et al*, 2009). The activating Ly49H receptor, encoded by *Klra8*, recognizes the MCMV-encoded m157 glycoprotein and leads to activation of Ly49H+ NK cells (Orr *et al*, 2009; Voigt *et al*, 2003). In humans, a similar activation of NK cells by human cytomegalovirus is observed with the preferential expansion of NKG2C+ NK cells (Guma *et al*, 2004; Hendricks *et al*, 2014; Ishiyama *et al*, 2022; Kim *et al*, 2019; Lopez-Verges *et al*, 2011). The expansion, contraction, and memory formation of Ly49H+ NK cells during MCMV infection bears similarity to antigen-specific responses of adaptive CD8+ T cells that generate memory CD8+ T cells (Adams *et al*, 2020; Buchholz *et al*, 2013; Kaech *et al*, 2002; Schlub *et al*., 2011).

A range of experiments investigated epigenetic changes, RNA transcription, and surface protein expression in single and bulk Ly49H+ NK cells undergoing expansion and contraction in mice responding to MCMV infection (Flommersfeld *et al*, 2021; Grassmann *et al*, 2019; Lau *et al*, 2018; Martinet *et al*, 2015; Min-Oo *et al*, 2014; Nabekura *et al*, 2014; Potempa *et al*, 2022; Riggan *et al*, 2022a). These observations show that the NK cell compartment undergoes phenotypic changes, resulting in changes in the abundances of varying subpopulations of cells. The development of these subpopulations depends on the three main signal inputs, antigen simulation of a specific receptor (e.g., Ly49H), co-stimulation, and cytokine signaling (Ni *et al*, 2012; Peng & Tian, 2017; Romee *et al*, 2016). These subpopulations express distinct epigenetic, transcriptomic, and protein markers imprinted by signaling and gene regulatory processes. Observations of clonal expansion of barcoded single Ly49H+ NK cells adoptively transferred into *Rag2^−/−^Il2rg^−/−^* mice revealed another layer of heterogeneity where sizes and compositions of NK cell clones originating from single naïve Ly49H+ NK cells vary widely (Flommersfeld *et al*., 2021; Grassmann *et al*., 2019). This heterogeneity in the NK cell population is regulated by the stochastic kinetics of cell proliferation, cell death, and differentiation involving the subpopulations that emerge as the NK cells respond to the infection. Analysis of similar heterogeneities in clonal CD8+ T cell populations (Buchholz *et al*., 2013; Gerlach *et al*, 2013) have shown generation of a highly proliferative short-lived effector phenotype and intrinsic stochastic fluctuations in the differentiation and proliferation processes, which account for heterogeneity of the CD8+ T cell clones (Buchholz *et al*., 2013). These effector CD8+ T cells are CD62L- and differentiate during the expansion phase from upstream memory precursors that are CD62L+(Schlub *et al*, 2010; Yang *et al*, 2011). More recently, tracking proliferation, death, and differentiation of CD8+ T cells during clonal expansion has introduced the notion of division destiny where antigen and cytokine stimulation in an individual T cell determines the division destiny or the number of divisions the T cell undergoes before differentiating into a state with negligible proliferation (Cheon *et al*, 2021; De Boer & Yates, 2023; Hawkins *et al*, 2007; Heinzel *et al*, 2017). More generally, these studies demonstrated that the fates of individual T or B cells during clonal expansion are linked to the number of divisions executed by those individual cells (Bresser *et al*, 2022; De Boer & Yates, 2023). Modeling the expansion phase of CD8+ T cell clones incorporating division destiny, heterogeneous antigen stimulation times, inheritance of division and death times in daughter cells from mother T cell, and cell division- dependent kinetics of cell lineage marker (e.g., CD62L) expressions (Pandit & De Boer, 2019) showed that the model can qualitatively recapitulate the observed heterogeneity (Buchholz *et al*., 2013) in the CD8+ T cell population in the expansion phase. The above models known as Cyton models investigate stochastic kinetics of individual T/B cells during clonal expansion and contraction. The existence of division destiny in antigen stimulated Ly49H+ NK cells still needs to be elucidated (Rückert & Romagnani, 2024).

Despite the qualitative similarities between the antigen-specific responses of NK cells and CD8+ T cells there are several marked differences. Single-cell NK clones typically undergo a relatively smaller 1000× increase in number compared to the 40,000× increase in CD8+ T cell populations (Adams *et al*., 2020). Additionally, mature CD27- NK cells grow more slowly than their immature CD27+ counterparts (Buchholz *et al*., 2013; Hayakawa & Smyth, 2006; Paul *et al*, 2016), whereas effector CD62L- CD8+ T cells proliferate faster than precursor CD62L+ subsets (Adams *et al*., 2020; Kretschmer *et al*, 2020). These data suggest there may be mechanistic differences in antigen-specific NK and CD8+ T cell responses.

Here, we addressed whether distinct mechanisms underlie the expansion of NK cells and CD8+ T cells during antigen-specific responses by performing stochastic population dynamic modeling on published bulk and single-cell NK cell data collected during the expansion phase following MCMV infection in C57BL/6 mice. Our analysis and modeling show that the mechanistic rules that describe the expansion of CD8+ T cells are inadequate for describing the development of MCMV-specific NK cells. We found that mature CD27-Ly6C- NK cells undergo a greater rate of cell proliferation compared to immature CD27+ NK cells and more mature CD27-Ly6C+ NK cells, and mature CD27- NK cells undergo increased death relative to the CD27+ immature cells. The variations in the timescales of proliferation and death processes coupled with the intrinsic stochastic fluctuations within these processes among the NK cell subsets regulate the heterogeneity in the sizes of the NK cell clones originated from individual immature NK cells during MCMV infection. Quantification of the population kinetics of Ly49H+ NK cells in MCMV infection in the expansion phase shows a 10-fold increase in the cell growth rate compared to that in homeostatic conditions, whereas the effect of the infection on the maturation rate appears to be negligible. We also find that growth rates of adoptive NK cells in transfer experiments during MCMV infection in the expansion phase are higher than those of endogenous NK cells.

## Results

### A minimal progressive two-state immature to mature differentiation model cannot capture the heterogeneity of the NK cell clones in the expansion phase

We began our investigations by studying whether a minimal two-state progressive differentiation model (Fig. 1a) could describe the size and composition of NK cell clones that originated from single immature Ly49H+ CD27+ NK cells at day 8 post-infection with MCMV, a time typically at the peak of expansion. The differentiation of immature CD27+ to mature CD27- NK cells during MCMV infection and in homeostasis has been widely documented and established in previous experiments (Chiossone *et al*, 2009; Grassmann *et al*., 2019; Min-Oo *et al*., 2014; Mitrovic *et al*, 2012). We used data collected by Flommersfeld *et al.(Flommersfeld et al., 2021)*, in which they adoptively transferred single genetic marker barcoded Ly49H+ NK cells to a host that is subsequently infected with MCMV. Splenocytes are then harvested, and resulting NK cell clones are distinguished with flow cytometry, and CD27+ and CD27- cells are determined with flow cytometry gates. We characterized the NK cell clones by the means and variances of the populations of CD27+ or CD27- NK cells and the co-variance between the two cell types in the clones. The NK cell clones reported by Flommersfeld *et al*. contained mixtures of CD27+ and CD27- NK cells in the spleen. We evaluated the percentage of CD27+ NK cells in each clone and computed the correlation (*C_size-CD27+_*) of the size of the clone with the percentage of CD27+ NK cells in the clones. The larger size NK cell clones contained a greater proportion of the CD27- NK cells and thus produced a negative correlation between the clone size and the percentage of CD27+ NK cells in individual clones (Fig. 1b). This finding is in line with that found in a similar experiment by Grassmann *et al. (Grassmann et al., 2019).* These authors also transferred congenically marked CD27- and CD27+ Ly49H+ NK cells in 1:1 ratio at day 0 and then assayed the expanded CD27+/- NK cells at day 8 post infection, showing a higher proportion of mature CD27- NK cells compared to the immature CD27+ NK cells (Figure S2E in Grassmann *et al. (Grassmann et al., 2019)*). This result is supported by other experiments (Hayakawa & Smyth, 2006; Paul *et al*., 2016), and we used this observation to constrain our model parameter space. Heterogeneity of clones of CD8+ T cells generated by antigenic SIINFEKL stimulation showed a similar pattern where the largest clones in the expansion phase (day 7 post-stimulation) are composed of higher fractions of the mature CD62L- CD8+ T cells (Buchholz *et al*., 2013). Because mature CD62L- CD8+ T cells proliferate faster than the immature CD62L+ CD8+ T cells (Buchholz *et al*., 2013; Kretschmer *et al*., 2020), it is intuitive that a larger size of a clone can be attributed to the larger proportion of the mature CD8+ T cells in the clone (Fig. 1c). In contrast, NK cells proliferate slower when they differentiate to the mature CD27- phenotype. Thus, it is unclear how the similar relationship between the clone size and the composition of the clones can be understood. To probe this question further we used a stochastic model (Fig. 1a) that describes the proliferation and differentiation processes in single Ly49H+ NK cells in response to MCMV infection. We do not model the MCMV infection explicitly and assume that the parameters of the model capture the effects of antigen, co-receptor, and cytokine stimulation and signaling and epigenetic modifications on the rates of differentiation and the proliferation of the cell types. In this model, immature CD27+ NK cells differentiate with a rate *r* to mature CD27- NK cells, and each cell type proliferates with rates, *k_I_* and *k_M,_* respectively. The proliferation and differentiation processes are modeled as stochastic events occurring with the above rates. We start our simulations with a single CD27+ NK cell and evaluated the number of CD27+ and CD27- NK cells at 8 days, which represent the size and the composition of a single NK cell clone. Repetitions of the stochastic simulations with the same initial configuration and rates will produce clones of different sizes and compositions due to intrinsic noise fluctuations in the stochastic simulations. We computed the correlation (*C*_size-CD27+_) between the clone sizes and the percentages of the CD27+ NK cells in each clone from our simulations. Our parameter scan of randomly sampled values of *k_I_*, *k_M_*, and *r* revealed that *C_size-CD27+_* is negative *only* when *k_I_*< *k_M_* (Fig. 1d). However, the observations that CD27+ cells should grow faster than CD27- cells imply *k_I_* > *k_M_* for the NK cells. Therefore, the two-state model with no death cannot capture the heterogeneity of the NK cell clones.

**Figure 1:**
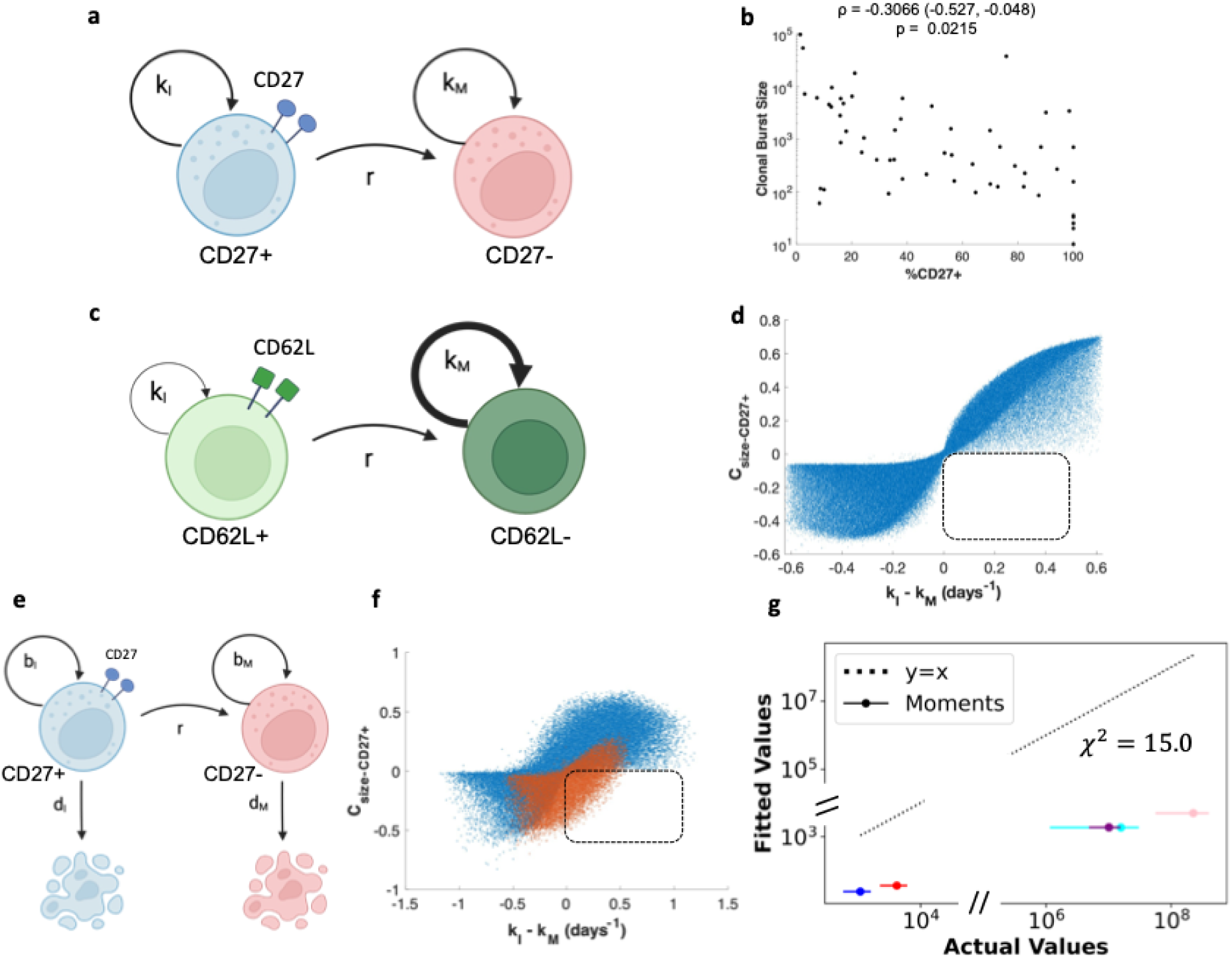
Two-stage linear models cannot capture desired properties of experimental clonal burst data. **(a)** Schematic representation of the two-stage model. The model describes immature (CD27+) cells and mature (CD27-) cells. Each cell type has a distinct growth rate (*k_I_* for immature, *k_M_* for mature), and immature cells differentiate to mature cells at rate *r*. (**b)** Experimental data from Flommersfeld *et al*. display negative value for the correlation *C_size-CD27+_* between clonal burst size and %CD27+ cells. **(c)** Schematic description of antigen specific clonal proliferation and differentiation in CD8+ T cells. Because mature CD62L- cells proliferate more rapidly compared to the immature CD62L+ cells, initial clones made primarily of CD62L- cells can grow to generate the larger size clones. **(d)** A parameter scan of all possible combinations of *k_I_*, *k_M_*, and *r* shows that negative correlations (*C_size-CD27+_* < 0) are not realized when *k_I_ > k_M_*. Each point represents a value of *C_size-CD27+_* obtained from the model at a unique value of set of model parameters. The parameter configurations that populate the bottom-right quadrant satisfy our two constraints imposed by experimental observations regarding the negative values of *C_size-CD27+_* and higher growth rate of immature NK cells compared to their mature counterparts. (**e)** Schematic representation of the three-state model including cell death. The model has separate parameters for birth of immature (CD27+) cells (*b_I_*), death of immature cells (*d_I_*), birth of mature (CD27-) cells (*b_M_*), death of mature cells (*d_M_*), and differentiation of immature cells to mature (*r*). In this model, the growth rates are defined as *k_I_=b_I_-d_I_* and *k_M_=b_M_-d_M_*. These net rates relate to the rates of growth or loss of clonal populations. **(f)** A parameter scan for this model shows that some parameter configurations can meet the two constraints where *k_I_ > k_M_* and the parameters result in negative values for *C_size-CD27+_*. Each point represents a unique parameter configuration. Red points denote parameter configurations where *b_M_ > b_I_* <u>and</u> *d_M_ > d_I_*. **(g)** Shows the best fit of the model in (e) to the moments with the constraints *k_I_ > k_M_* and *C_size-CD27+_* < -0.2. Circles represent the fitted values of the first and second moments for the distribution of the NK cell clones obtained from the model. Blue and red circles represent mean populations of CD27+, and CD27- NK cells, respectively. The variances of the CD27+ and CD27- NK cells are shown in cyan and pink circles, and the covariance between the CD27+ and CD27- NK cells is shown in purple. Horizontal lines around the circles show the standard deviations of bootstrapped moments. The solid line is the line *y=x*, which demonstrates where values would lie if we had a perfect fit.

Next, we investigated if inclusion of death of NK cells can make the model consistent with the negative correlation between the clone size and the percentage of CD27+ NK cells in each clone. We updated our proposed model to include both proliferation and death of each cell population (Fig. 1e). In such a model, each cell has a probability of proliferating according to birth rate *b* and probability of dying indicated by the death rate *d*. Cells can go through many proliferation events before undergoing death, which would indicate a positive net growth rate for the clone. The net growth of the CD27+ and CD27- NK cell populations are determined by *k_I_=b_I_-d_I_* and *k_M_=b_M_-d_M_*, respectively, where *b_I_* and *d_I_* (or *b_M_* and *d_M_*) are the proliferation and death rates of the immature CD27+ (or mature CD27-) NK cells. Repeating a parameter scan by randomly sampling the 5 parameters (*b_I_, d_I_, b_M_, d_M_,* and *r*) from uniform distributions and evaluating the correlation *C_size-CD27+_* shows that some parameter configurations can generate the negative values at the parameter values, *k_I_*> *k_M_* (Fig. 1f). Further inspection shows that parameter values in the regime *k_I_* > *k_M_* that produce the negative correlation share a common trait: mature CD27- cells proliferate *and* die faster than immature CD27+ cells. To gain further insight into this behavior we investigated the dependence of *C_size-CD27+_* on the parameter values and stochastic fluctuations analytically and found that larger proliferation and death rates of the CD27- NK cells produce large stochastic fluctuations in the sizes of the CD27- cell populations. Thus, mature CD27- NK cells in a clone could undergo many proliferation events before encountering death, creating larger clones with higher proportion of mature CD27- NK cells. Conversely, the CD27- cells could go through many death events before a proliferation event causing the clone size to shrink (details in Supplementary Text 1).

We investigated whether this model could accurately fit the first (means) and second (variances and covariance) moments of the measured CD27+ and CD27- NK cell populations at 8 days post-infection. We fit the model by minimizing the sum of square errors (SSE) subject to the constraints *C_size-CD27+_* < -0.2 and *k_I_ > k_M_*. This parameter configuration cannot accurately capture the moments of the clonal burst data (Fig. 1g). Thus, although the model that incorporates both NK cell proliferation and death is able produce the negative correlation between the clone size and the percentage of CD27+ NK cells in each clone, it poorly describes the distributions of the immature and mature NK cells in the clones.

To further investigate the role of NK cell death in explaining the clone size distribution we compared the above two-state models without or with NK cell death based on the theory of model selection using modified Akaike Information Criterion (AIC_c_) (Burnham *et al*, 2011). We computed the probability distribution function of the number of immature CD27+ (N_I_) and mature CD27- (N_M_) cells at any time obtained from our stochastic model by solving the corresponding Master equation numerically and computed the Likelihood function for the available clone size distribution data (details in the Materials and Methods section), and then evaluated modified-AIC (AIC_c_) for the two models. To keep the computation in a feasible range we used a part (38 out of 56 clone sizes) of the available data where the maximum clone size is ∼ 1000. The subset of clone size data used here also shows a negative value of *C_size-CD27+_*. The AICc value for the two-state model with death is over 50 points lower than that for the two-state model without death, which clearly show that the data favor the model with NK cell death. Although this approach only models a subset of the data, it also finds that birth and death rates for mature cells are greater than that of immature cells, and that the net growth rate of the clones of immature cells is greater than that of mature cells. Further details regarding the analysis are provided in the Materials and Methods section and the Supplementary Material. Because of the computationally intensive nature of this approach and its agreement with the simpler method of fitting moments with a constraint, we continue with the later approach for subsequent models. In addition, we investigated the roles of features considered in Cyton models for T cells (Pandit & De Boer, 2019) such as heterogeneity in the duration of antigen stimulation (Fig. S2), and the dependence of cell differentiation rates on the number of cell division (Fig. S3). The model where the durations are distributed in a log-normal distribution, as well the model where the rate of differentiation increased linearly with the number of cell divisions, both generated negative values of *C_size-CD27+_* when *k_I_* > *k_M_*, without cell death (Fig. S2b-d, S3b). Because the distribution of the duration of activation of NK cells in CMV infections as well as the increasing rate of differentiation with increasing number of cell divisions are not established for NK cells, we did not move forward with fitting these models to clonal burst moments. We also investigated an NK cell proliferation model where the immature NK cell divides asymmetrically to generate a mature and an immature NK cell (Fig. S4). T cells can undergo asymmetric division where the daughter cells proximal and distal to the antigen-presenting cell are predisposed toward the effector and the memory fates, respectively (Chang *et al*, 2007); though asymmetric division has not been observed in NK cells. This model generated negative values of *C_size-CD27+_* but did not provide good agreement with the moments. The above alternative models contain more parameters as well as additional mechanisms that were not validated in experiments better than the minimal model we considered here. However, these explorations suggest that additional features needed to be included in the minimal model to capture the heterogeneities observed in clonally expanded Ly49H+ NK cells. The kinetics of clonal populations arising from these processes can be described by ordinary differential equations which do not permit aggregation of division and death rates, consequently, parameter estimation requires independent measuremets of proliferation (e.g., Ki67) or death. In the next sections, we study some extensions of our two-state model that can be rationalized against available experiments.

### A progressive three-stage differentiation model can quantitatively recreate clonal burst dynamics

We modified our two-state model to include an intermediate state (Fig. 2a) in the model. This reflects many investigations probing NK cell maturation, which show NK cell development goes through several maturation states and some of those states can potentially represent the proposed intermediate state in the model. A well-established NK cell maturation scheme is CD27+CD11b- ➔ CD27+CD11b+ ➔ CD27-CD11b+ (Chiossone *et al*., 2009). However, the clonal burst data show a negligible proportion of CD11b- NK cells at 8 days post-infection. Another potential maturation scheme could be CD27+KLRG1- ➔ CD27+KLRG1+ ➔ CD27-KLRG1+ (Kamimura & Lanier, 2015). In addition, recent single cell RNA-seq experiments show the presence of CD27-Ly6C+ and CD27-Ly6C- cells at 8 days post-MCMV infection (Riggan *et al*., 2022a), thus the intermediate state could represent a CD27-Ly6C- state as well. Here we present results for the model where the intermediate state represents a mature CD27- state, the other variations of the model are shown in the Supplementary Material (Fig. S5-S7). To determine parameter ranges that give rise to higher growth of mature NK cells compared to their immature counterparts, we computed the average size of a clone that originates from either a single CD27+ NK cell or single CD27- NK cell by solving ODEs describing the kinetics of the average NK cell populations. We constructed an effective growth difference metric Δ*_k_,* which describes the difference between the average NK cell populations at day 8 that originate from a single CD27+ or CD27- NK cell at t=0. If Δ*_k_* is positive, then CD27+ cells grow faster than CD27- cells (Materials and Methods). We performed a parameter scan on this three-stage model and find that some parameter configurations with Δ*_k_* > 0 can capture the negative values of *C*_size-CD27+_ (Fig. 2b). We then evaluated whether this three-stage model can accurately describe the distributions of the immature and mature NK cells in clones. We fitted the means, variances, and the covariance of the NK subpopulations in the clones by varying the model parameters with constraints Δ*_k_* > 0 and *C*_size-CD27+_ < -0.2 and found that the best-fit three-stage model shows an excellent fit to the data (Fig. 2c, Table 1). This best-fit parameter set generates a *C*_size-CD27+_ of -0.351, well within the confidence bounds of the correlation coefficient of the data (Fig. 2d). Thus, adding an intermediate CD27- stage to the model can capture all the key features of the heterogeneity in the NK cell clonal populations.

**Figure 2:**
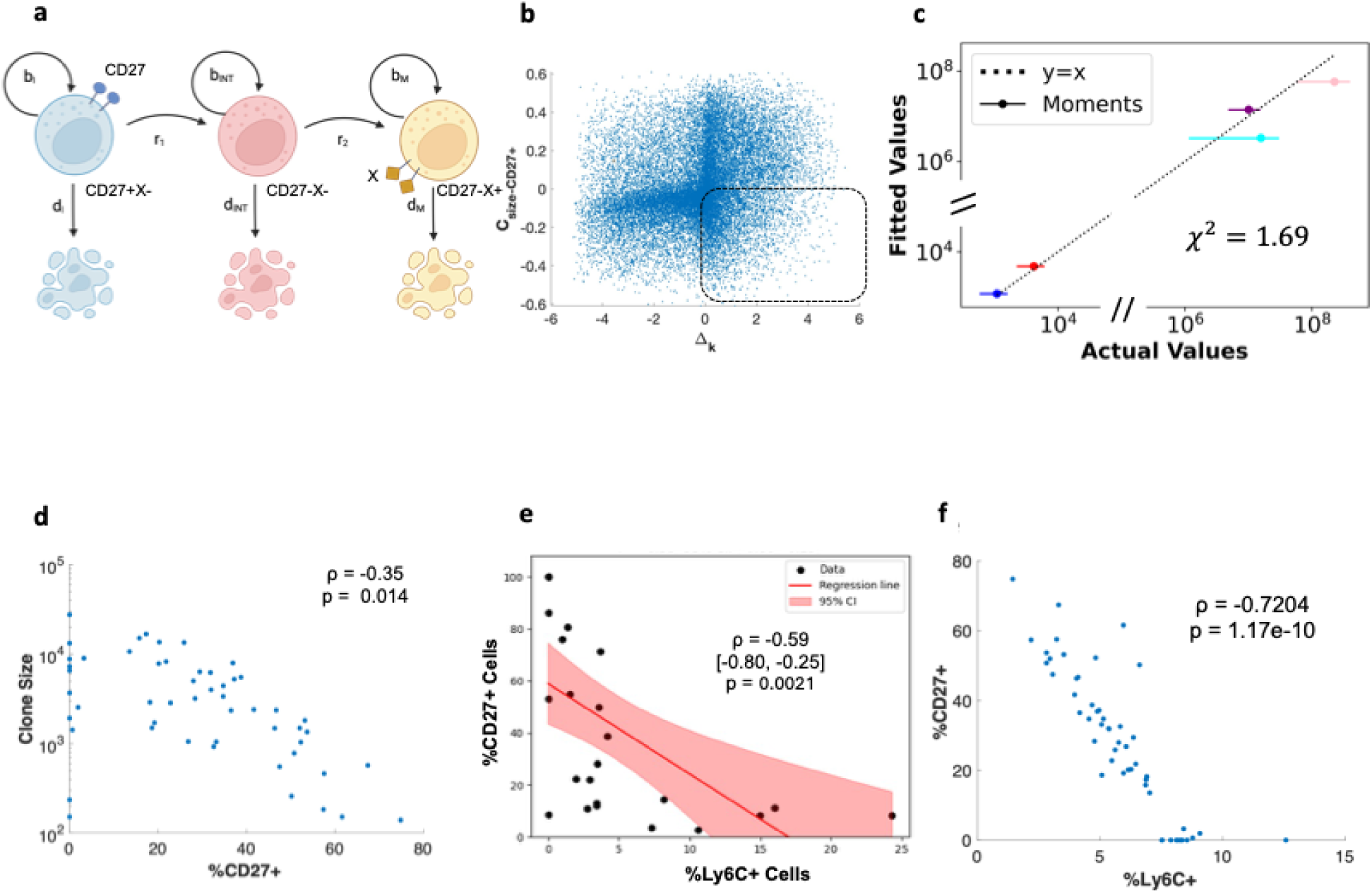
A three-stage linear model can capture experimental observations of NK cell clones if intermediate double-positive NK cells are the fastest growing subset. **(a)** Schematic representation of the three-stage model. In this model there are three stages of NK cell maturation: an immature (CD27+X-), intermediate (CD27-X-), and terminally mature (CD27- X+) subset, where the terminally mature state is marked by an expression of a yet to be determined marker protein X. Each subset has a distinct proliferation and death rate, and cells progress from immature to intermediate maturity according to rate *r_1_* and from intermediate to terminal maturity according to rate *r_2_*. (**b)** A parameter scan for the model in (a) shows that adding a third stage of maturation can account for the negative values in *C_size-CD27+_*. Each point represents a unique parameter configuration. Positive values of Δ*_k_* indicate that immature CD27+ cells grow faster than mature CD27- cells and vice versa. See the Materials and Methods for further details. **(c)** Best fit of this model to the moments with the constraints Δ*_k_* > 0 and *C_size-CD27+_* < -0.2. Circles represent the fitted values of the moments to the model. Blue and red circles represent mean populations of CD27+, and CD27- NK cells, respectively. The variances of the CD27+ and CD27- NK cells are shown in cyan and pink circles, and the covariance between the CD27+ and CD27- NK cells is shown in purple. Horizontal lines around the circles show the standard deviations of bootstrapped moments. The solid line is the line *y=x*, which demonstrates where values would lie if we had a perfect fit. **(d)** Clones stochastically simulated with Gillespie’s algorithm (Materials and Methods) from the best fit parameters and the resulting clone sizes, CD27+ percentages, and correlation. **(e)** Observed clonal compositions of Ly6C+ and CD27+ cells from stochastically simulated clones. **(f)** Simulated clones from the three-stage model in (a) at the best fit parameter values shown in (c) when the three stages are taken as immature CD27+Ly6C-, mature CD27-LyC- and the terminally mature CD27-Ly6C+, respectively. Each point is a simulated clone resulting from a single NK cell.

**Table 1:**
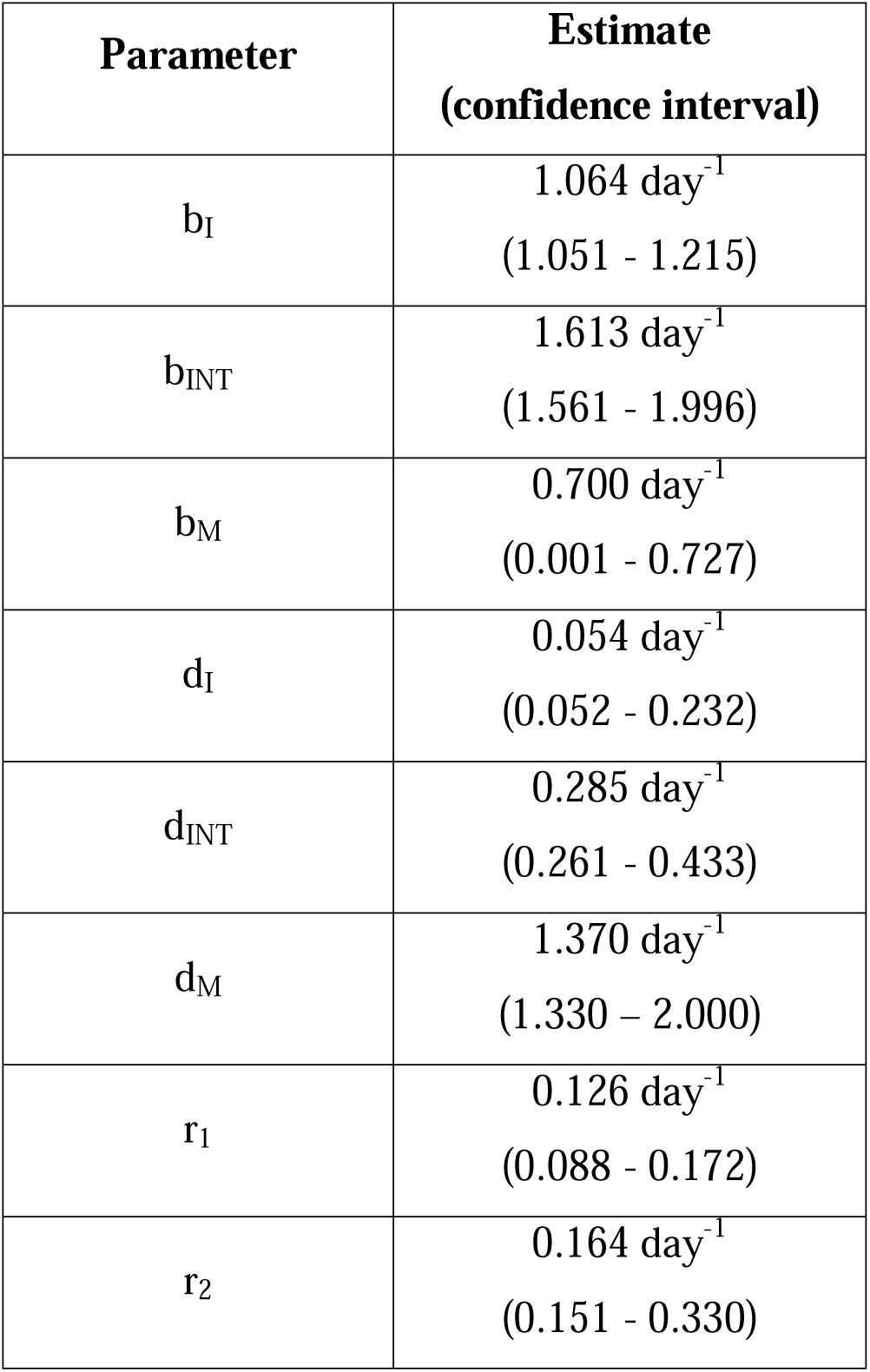
Best fit parameters to 3-stage model describing NK clonal bursts given in Fig. 2a. Confidence intervals are determined by bootstrapping.

| Parameter | Estimate<br>(confidence interval) |
| --- | --- |
| $b_I$ | 1.064 day<sup>-1</sup><br>(1.051 - 1.215) |
| $b_{\text{INT}}$ | 1.613 day<sup>-1</sup><br>(1.561 - 1.996) |
| $b_M$ | 0.700 day<sup>-1</sup><br>(0.001 - 0.727) |
| $d_I$ | 0.054 day<sup>-1</sup><br>(0.052 - 0.232) |
| $d_{\text{INT}}$ | 0.285 day<sup>-1</sup><br>(0.261 - 0.433) |
| $d_M$ | 1.370 day<sup>-1</sup><br>(1.330 - 2.000) |
| $r_1$ | 0.126 day<sup>-1</sup><br>(0.088 - 0.172) |
| $r_2$ | 0.164 day<sup>-1</sup><br>(0.151 - 0.330) |

The three-stage model above makes no assumptions about the identities of the differing phenotypes of the CD27- cell subsets. However, the best fit parameters indicate that the first CD27- subset has a high proliferation rate, and the final CD27- subset has a high death rate. Riggan *et al*. (Riggan *et al*., 2022a) previously determined that, in the contraction phase, Ly6C+ NK cells differentiate from Ly6C- cells, and that Ly6C+ cells die more rapidly than Ly6C- cells. Our model shows that similar activity may be observed during the expansion phase as well, as we show higher death of Ly6C+ cells and a unidirectional maturation from Ly6C- → Ly6C+ cells. Because our analysis pertains to the expansion phase of the NK cell response to MCMV we thus aimed to validate if the mature states proposed in the model can be identified with CD27-Ly6C- and CD27-Ly6C+ NK cells. We found that our best fit parameters correctly recreate the experimentally observed strong negative correlation between the %CD27+ and %Ly6C+ NK cells across clones (Grassmann *et al*., 2019), and correctly estimate that most clones have <20% Ly6C+ cells at day 8 post-infection (Fig. 2e-f). This independent validation gives evidence that our parameter estimates can be meaningfully interpreted for a model where NK cells differentiate along a trajectory from CD27+Ly6C- → CD27-Ly6C- → CD27-Ly6C+.

We sought to understand the mechanism by which the three-stage model can generate negative values for the correlation *C*_size-CD27+_ as well as quantitatively describe the heterogeneity in the NK cell populations in the clones. The two-stage model predictions for the first and the second moments for the NK cell populations are substantially less than the actual values, indicating that our best-fit parameters are not producing enough NK cells (Fig. 1g). The presence of the intermediate state in the three-stage model alleviates this issue, which can rapidly grow and increase the number of CD27-Ly6C+ NK cells while the rapidly dying CD27-Ly6C- NK cells can help meet the constraint that a population of CD27- cells grows slower than one of CD27+ cells. This variation increases stochastic fluctuations in the observed numbers of CD27- cells, as some clones will have many CD27- cells from high growth of the intermediate Ly6C-subset, while other clones will have few CD27- cells from low growth of the Ly6C+ subset. Furthermore, the presence of the CD27-Ly6C+ NK cells expand the range of the model parameters that can give rise to negative values of *C*_size-CD27+_.

### Comparison of kinetics between endogenous NK cells, adoptively transferred cells, and homeostatic cells identifies potential identical maturation rates in all conditions

Here we compare the kinetics of NK cell maturation occurring in response to MCMV infection and in homeostasis. In particular, we investigated the changes in the rates of cell growth and differentiation induced by MCMV infection. First, we quantified the growth and the differentiation rates using time-stamped flow cytometry data on homeostatic NK cells in peripheral blood with a one-time treatment of tamoxifen in fluorescent NK cell reporter mice reported by Adams *et al*.(Adams *et al*, 2021). Treating the mice with tamoxifen determines whether observed NK cells are newly populated from bone marrow or existed in peripheral blood at the onset of the infection. We fitted a two-state CD27+ → CD27- maturation model described by ordinary differential equations (see Materials and Methods) to the data collected at days 3, 7, 14, 23, 29, and 35 to estimate the growth rates of the immature CD27+ (*k_CD27_*_+_) and the mature CD27- (*k_CD27-_*) NK cells, the rate of differentiation (*r*) of the CD27+ to CD27- NK cells, and the rate (λ) with which CD27+ NK cells are populated from bone marrow. We estimated these rates both for Ly49H+ and Ly49H- NK cells. The estimated rates for Ly49H+ cells (Table 2, Fig. 3a) show that a CD27+ cell population would double every 6.9 days if there were no differentiation, whereas CD27- cells have a negative growth rate with a half-life of 5.6 days. This implies that the death rate of a population of CD27- cells is greater than their proliferation rate. The differentiation rate from CD27+ → CD27- phenotypes is similar to that of the death rate of Ly49H+ CD27- cells. Approximately 1.37% of the Ly49H+ NK pool is replenished from bone marrow every day. Comparing these rates to those of the Ly49H- NK pool reveals that Ly49H- NK cells are repopulated from bone marrow at almost twice the rate (∼2.4%/day). All other estimated rates for Ly49H- NK cells are within the confidence bounds of the Ly49H+ estimated rates (Table 2, Fig. 3b). Given the constraint governing λ which dictates

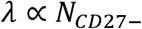

**Figure 3:**
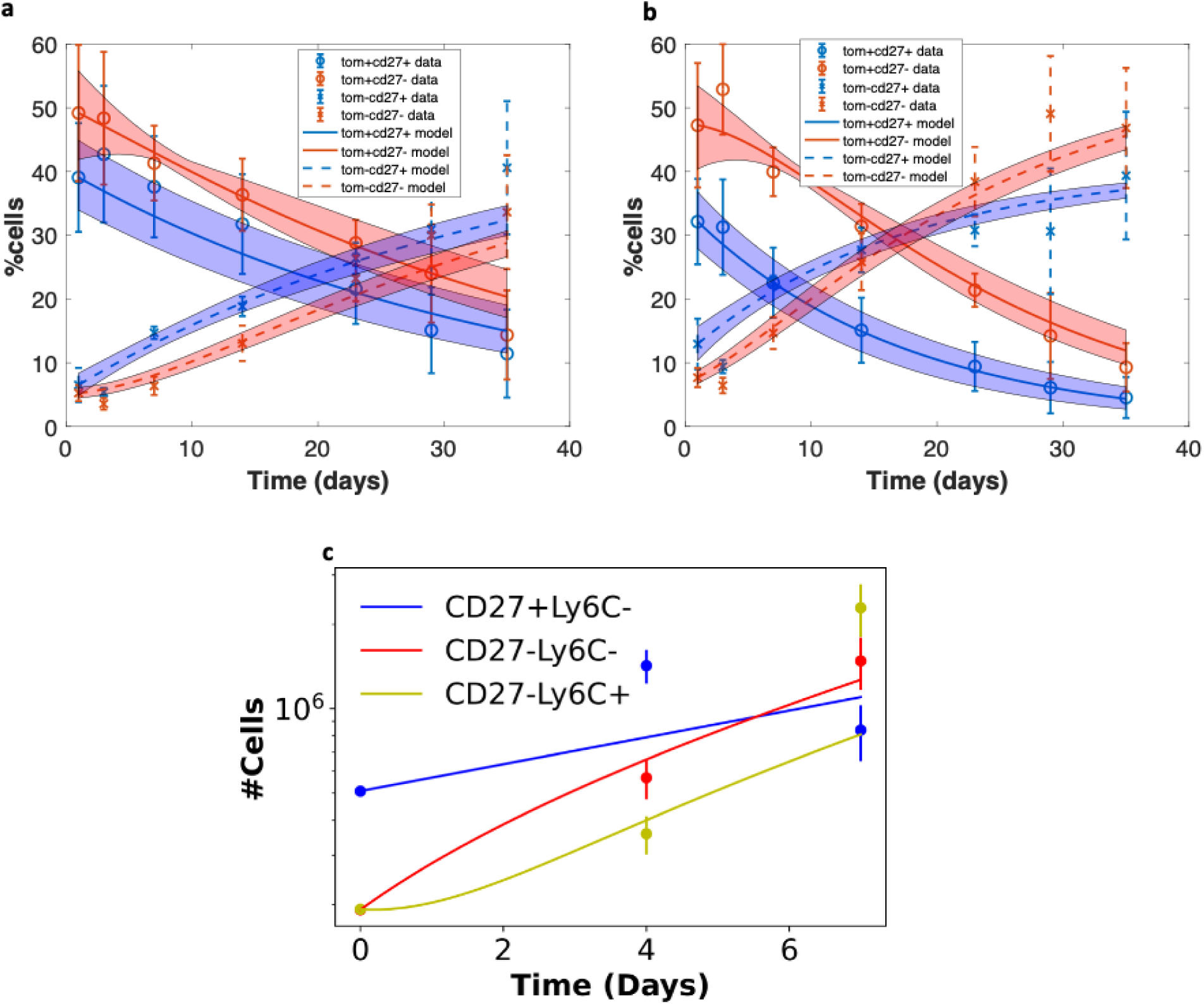
Kinetics of homeostatic NK cells and endogenous NK cells responding to MCMV infection. **(a-b)** Best fit to **(a)** Ly49H+ and **(b)** Ly49H- homeostatic NK cells. Y axis refers to percentage of the total Ly49H+ or Ly49H- population occupied by that NK subset. Circles and Xs represent the mean values of tamoxifen-induced td-Tomato positive and negative NK cells observed in the data, respectively. Solid and dashed error bars represent standard deviations around the means of tamoxifen-induced positive and negative NK abundances respectively. Smooth curves represent model fits for tamoxifen-induced positive (solid) and negative (dashed) NK cells. Shaded regions refer to model 95% confidence bands, as determined by bootstrapping model fits. (**c**) Best fit to profiled endogenous NK cells responding to MCMV infection. Points represent mean cell abundances, and error bars represent standard deviations of bootstrapped means. Smooth curves represent model fit.

**Table 2:** Best fit parameters to homeostatic Ly49H+ and Ly49H- NK cells. Confidence intervals are determined by bootstrapping.

| Parameter | Ly49H<sup>+</sup> Estimate<br>(confidence interval) | Ly49H<sup>-</sup> Estimate<br>(confidence interval) |
| --- | --- | --- |
| $\lambda$ | 1.366 percent/day<br>(1.117 – 1.594) | 2.412 percent/day<br>(2.028 – 2.915) |
| $r$ | 0.125 day<sup>-1</sup><br>(0.095 – 0.159) | 0.147 day<sup>-1</sup><br>(0.118 – 0.173) |
| $k_{CD27+}$ | 0.097 day<sup>-1</sup><br>(0.064 – 0.135) | 0.088 day<sup>-1</sup><br>(0.053 – 0.119) |
| $k_{CD27-}$ | -0.120 day<sup>-1</sup><br>(-0.152 – -0.091) | -0.107 day<sup>-1</sup><br>(-0.126 – -0.086) |

(Materials and Methods) the difference in estimated λ values can be attributed to a higher percentage of CD27- cells. Next, we compared these rates with corresponding rates estimated in our three-state model using the clonal burst data. The estimate for CD27+ → CD27- differentiation rate is equivalent between Ly49H+ NK cells in homeostasis and in MCMV infection. Our estimations for the clonal burst data (Table 1) show that varying subsets of CD27- cells have varying growth rates ranging -0.7 day^-1^ to 1.3 day^-1^in MCMV infection, a wide range not in disagreement with the estimate for homeostatic data. Ly49H+ CD27+ cells grow ∼10× faster in MCMV infection than in homeostatic conditions.

We investigated whether the kinetics of NK cell maturation in the expansion phase during MCMV infection in single-cell adoptive transfer experiments is quantitatively similar to that of an endogenous population of NK cells. To this end, we collected time-stamped mass cytometry data on C57BL/6 mice splenocytes at days 0, 4, and 7 post-MCMV infection (Potempa *et al*., 2022). We then gated Ly49H+ NK cells from each mouse and determined the mean relative abundances across mice of CD27+Ly6C-, CD27-Ly6C-, and CD27-Ly6C+ subsets among the NK cell population (Fig. S8). We multiplied these relative abundances by mean absolute numbers of Ly49H+ endogenous NK cells during MCMV infection from Robbins *et al*.(Robbins *et al*., 2004) to obtain time-stamped population sizes of NK cell subsets. We then modeled these data with the ordinary differential equation (ODE) solution in Equations S17-S25 using the parameters from Table 1 and an initial condition of the mean uninfected *in vivo* abundances of endogenous NK cells. The ODE result is multiple orders of magnitude above the measured abundances (Fig. S9a). This indicates that the kinetics of endogenous NK cells differ significantly from that of NK cells in adoptive transfer experiments. Further indication that the kinetics differ is the expansion of cells in endogenous conditions over 7 days is less than a 10- fold increase, while clonal expansion accounts for >1000-fold increase.

To further quantify the differences in the kinetic rates between the adoptively transferred and endogenous NK cells, we then fit the 3-stage model with 2 mature stages to the endogenous cell population. We used the gated Ly49H+ populations of CD27+/Ly6C-, CD27-/Ly6C-, and CD27-/Ly6C+ NK cells and fit to a system of ODEs that describe the net growth rates of each population and the differentiation rates between them (Materials and Methods). The best fit to this data has unrealistic estimates for the death rate of CD27-/Ly6C+ NK cells and the differentiation rate from CD27+ → CD27- cells (Table S5, Fig. S9b). We then attempted to constrain the fitting procedure by setting the CD27+ → CD27- differentiation rate to those of the previous estimations for the clonal burst kinetics and homeostatic conditions (Tables 1 and 2). Under this constraint, the only rate estimated to be within the confidence interval for the equivalent rate in adoptive transfer experiments is the differentiation rate from Ly6C- → Ly6C+ (Table 3, Fig. 3c). The growth rate of CD27+Ly6C- cells is ∼4× faster for adoptively transferred cells, the growth rate of CD27-Ly6C- cells is 2-3× faster, and the death rate of mature cells is ∼1.5× faster. However, the resulting best fit to the endogenous NK cell population does show that the intermediate CD27-/Ly6C- cell population has the fastest growth rate, and the mature CD27-/Ly6C+ cells have a large death rate, confirming the qualitative results from the adoptive transfer experiments. Thus, although the growth kinetics are different between endogenous and adoptive transfer experiments, both systems have the intermediate population proliferating rapidly and the terminally mature population dying rapidly.

**Table 3:** Best fit parameters to endogenous NK cells responding to MCMV infection with constrained *r_1_*. Confidence intervals are determined by bootstrapping.

| Parameter | Estimate<br>(confidence interval) |
| --- | --- |
| $k_I$ | 0.236 day<sup>-1</sup><br>(0.153 - 0.350) |
| $k_{INT}$ | 0.537 day<sup>-1</sup><br>(0.065 - 0.874) |
| $k_M$ | -0.484 day<sup>-1</sup><br>(-1.194 - 0.371) |
| $r_1$ | 0.126 day<sup>-1</sup><br>(fixed) |
| $r_2$ | 0.447 day<sup>-1</sup><br>(0.000 - 0.874) |

Both the adoptively transferred cells and endogenous population are estimated to have a large death rate for CD27- cells. To test this result, we analyzed NK cell viability in mice that were infected with MCMV at different times post-infection (Fig. 4a). Surprisingly, mature (CD27-) cells indeed had a greater fraction of dead cells than immature (CD27+) cells during expansion (days 4 and 7 post-infection) (Fig. 4b). To determine if the higher rates of cell death in the CD27- NK cell population in the three-state model give rise to a higher fraction of dead NK cells we extended our three-state model to evaluate the number of dead mature and immature NK cells (Fig. S10c) which showed that the best fit parameters for the adoptive transfer experiments are consistent with the results reported above (Fig. 4c). Because we only have best fit growth rates of the endogenous NK cells and not separate proliferation and death rates, we tested a variety of birth and death rates consistent with the best fit parameters such that *b-d=k* for each cell subset. Many of the possible configurations agree with the higher percentage of live CD27+ cells. This gives additional confirmation that CD27- cells may in fact be dying at faster rates than CD27+ cells.

**Figure 4:**
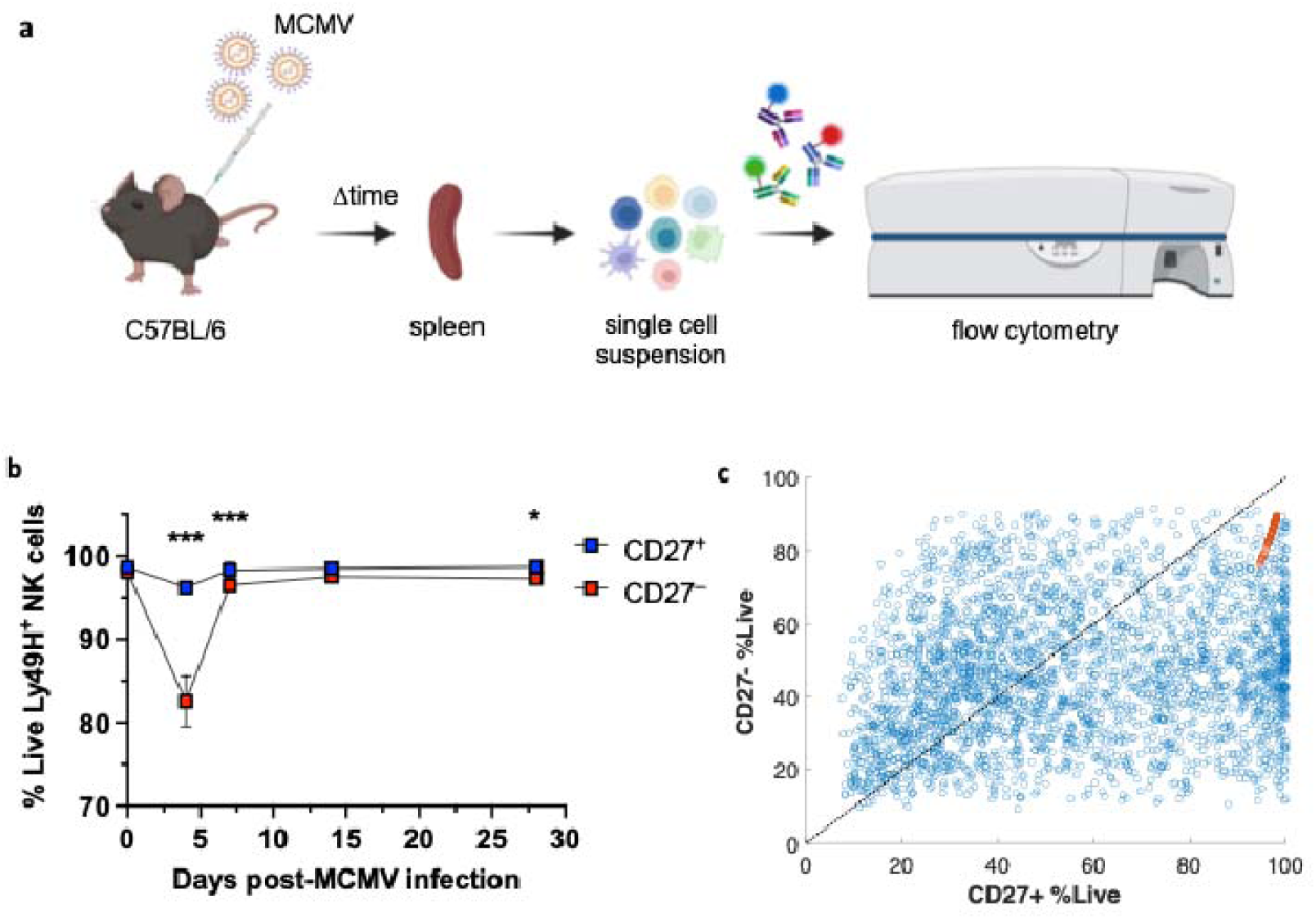
Mature CD27- NK cells undergo rapid death during the expansion phase. **(a)** Schematic of experimental setup. **(b)** Relative abundances of dead cells for CD27+ (blue) and CD27- (red) *ex vivo* NK cell subsets as measured by flow cytometry experiment. Data are representative of experiments with n = 4-5 mice per timepoint. Graph shows means ± SEM; *P <0.033, and ***P <0.001 represent statistically significant difference between CD27^+^ and CD27^-^ NK cells as determined by 2-way ANOVA. **(c)** Parameter scans for determining percentage of live cells for CD27+ and CD27- subsets using the model shown in Fig. S10c. Blue points represent a parameter scan where birth and death rates were varied by random sampling such that the net growth rates k_I_ = b_I_-d_I_, k_INT_ = b_INT_-d_INT_, and k_M_ = b_M_-d_M_ are equivalent to the kinetic estimates for endogenous NK cells shown in Table 3. Red points represent randomly sampled dead cell clearance rates with other rates set to those describing adoptive transfer kinetics in Table 1. Black dotted line is *y=x*, and points below this line show a higher percentage of live cells for immature CD27+ cells than for mature CD27- cells.

## Discussion

We employed stochastic minimal mechanistic *in silico* models to investigate mechanisms that underlie development of MCMV-specific NK cells at early times (0 – 8 days post-infection). We found Ly49H+ NK cells undergoing antigen-specific expansion follow a maturation scheme from (immature) CD27+Ly6C- → (mature I) CD27-Ly6C- → (mature II) CD27-Ly6C+, characterized by a highly proliferative mature CD27-Ly6C- phenotype and a more mature CD27- Ly6C+ phenotype with higher rates of cell death (Fig. 5). The differences in the rates of proliferation and cell death are crucial to give rise to the heterogeneous clones that originate from single immature CD27+ NK cells, in particular, the presence of larger sized NK cell clones composed of mature CD27- NK cells at 8 days post-infection. Similar heterogeneity in the clones of CD8+ T cells generated in the expansion phase (<8 days post-infection) from an acute antigen-specific expansion of single immature CD62L+ cells in C57BL/6 mice has been observed (Buchholz *et al*., 2013); however, the larger CD8+ T cell clones composed of effector CD62L- arise from the higher rate of proliferation of the CD62L- CD8+ T cells. This is in contrast with the NK cell maturation program where the larger sized clones of mature NK cell populations arise due to the interplay of uneven increase in the proliferation rates and increase in the rates of cell death as the NK cells mature, and intrinsic stochastic fluctuations occurring in these processes. Innate heterogeneity in adoptively transferred single NK cell phenotypes may also contribute to observed stochasticity. In CD8+ T cells, precursor CD62L+ cells are deemed memory precursors, matching their lowly proliferative phenotype (Kretschmer *et al*., 2020). An equivalent population of memory precursor NK cells has not been identified. CD27-Ly6C+ cells match previously defined phenotypes for memory NK cells and could potentially be memory precursors. However, our model shows that these cells experience rapid cell death. Therefore, for these cells to live longer to become memory cells additional processes such as a change in environmental conditions or an epigenetic change that promotes survival in these cells at later times during the contraction phase could become relevant. This differentiation trajectory is also in contrast with CD8+ T cells, as NK cell memory precursors would be formed downstream of effector (Ly6C-) NK cells, whereas CD8+ effectors (CD62L-) are formed downstream of CD8+ T cell memory precursors (CD62L+) in acute infection. Alternately, another mature NK subset such as Ly6C-KLRG1- that is lowly proliferative and long-lived could give rise to the adaptive NK cell phenotype. Inclusion of these processes as well as the later time kinetics could be a future extension of our approach. We note a similarity of our three state NK cell maturation model with maturation of T cells in persistent (13-140 days) *T. gondi* infection in genetically resistant mice reported by Robey and colleagues (Chu *et al*, 2016) where the infection gave rise to a continuous production of CXCR3-KLRG1+ terminally differentiated effector T cells from a quiescent memory CXCR3+KLRG1- T cells via an intermediate CXCR3+KLRG1+ state. The intermediate T cell state showed higher proliferation and increased death compared to the memory and effector T cells. However, these T cells were uniformly CD62L- in contrast to the memory and effector T cells that are produced in response to acute antigenic SIINFEKL infection.

**Figure 5:**
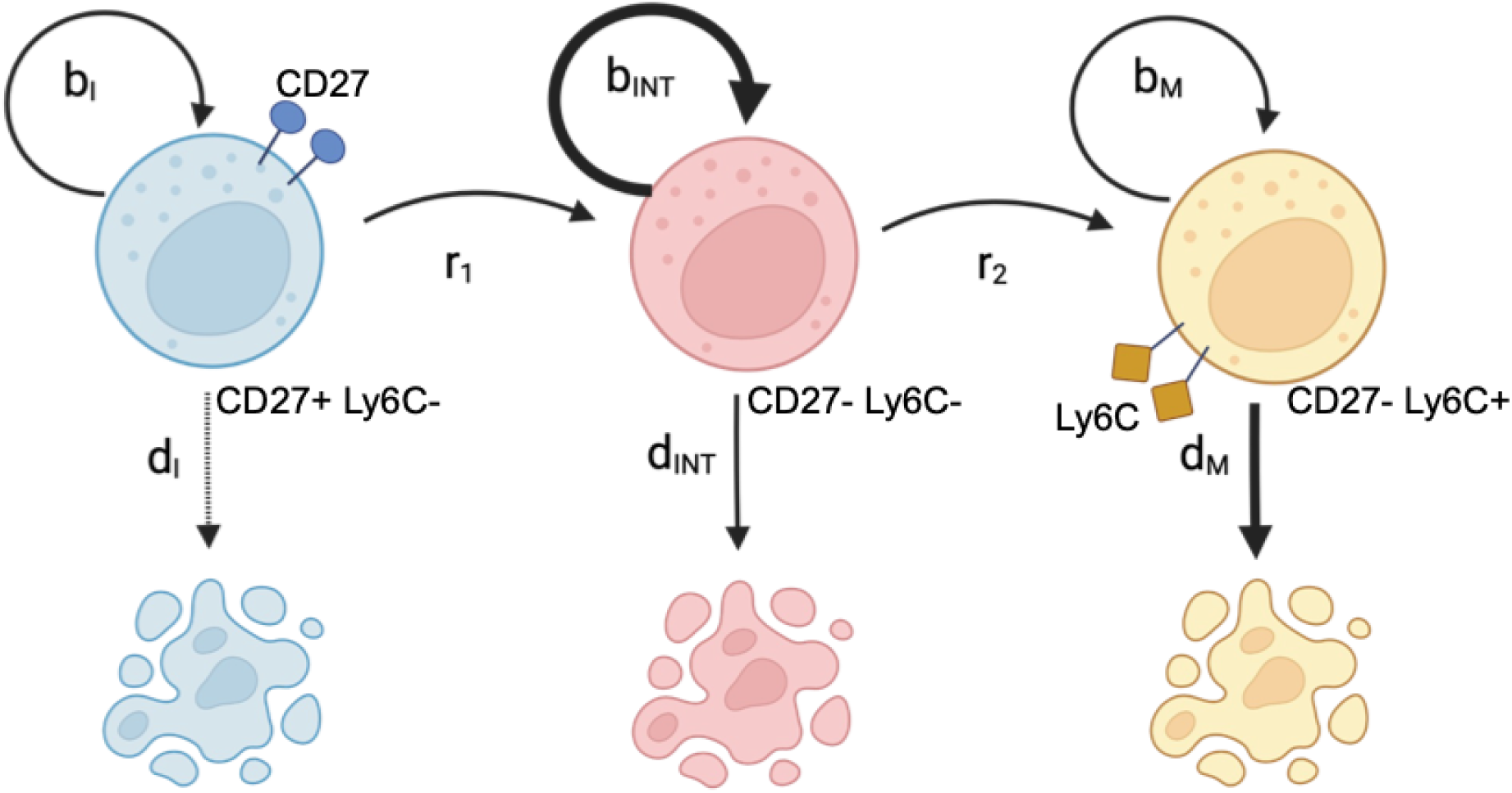
Proposed mechanism of NK clonal expansion in response to MCMV infection. Three distinct subsets of NK cells are necessary to generate full NK response to infection. The first is CD27+, Ly6C-, and moderately proliferative, with a doubling time of ∼0.3 days and negligible cell death. The second is CD27-, Ly6C-, and highly proliferative, with a doubling time of ∼0.2 days. The last is CD27-, Ly6C-, and prone to rapid cell death with a half-life of ∼0.45 days. This picture paints CD27-Ly6C- NK cells as having a highly proliferative and highly effective phenotype, similar to effector CD8+ T cells, and CD27+Ly6C- cells as long-lived, similar to memory precursor CD8+ T cells.

Integration of signals in NK cells during MCMV infection leads to epigenetic remodeling affecting NK cell differentiation, proliferation, and death (Riggan *et al*, 2022b; Rückert & Romagnani, 2024; Santosa *et al*, 2023). In T cells, the sum of signals received during clonal expansion has been found to regulate the number of division and the time of death or the division destiny in individual T cells that can arise due to a global remodeling of the transcriptional and epigenetic program (Iwata *et al*, 2017; Rückert & Romagnani, 2024). We considered a variant of the Cyton models (De Boer & Yates, 2023), which has been employed in modeling clonal expansion in T cells, where the rate of differentiation from the immature to the mature state depends on the number of divisions in individual NK cells. This model can give rise to negative values in the correlation between the clone size and the percentage of immature CD27+ NK cells in the absence of NK cell death, suggesting that inheritance of epigenetic states regulating cell fates in individual NK cells can play an important role in clonal expansion. Furthermore, we also find the presence of different time delays in NK cells to encounter appropriate activation signal in a pure birth model with two states can give rise to negative values in *C*_size-CD27+_ implicating a role of different activation times during the clonal expansion. The exploration of division destiny and activation times in NK cells during clonal expansion could be an exciting future direction.

NK cell clones in MCMV infection go through a smaller (∼1000×) fold increase in the expansion phase compared that (∼40,000×) of antigen-specific expansion of CD8+ T cells. This could be because CD8+ T cells can generate autocrine cytokines such as IL-2 that help the cells to proliferate (Kalia & Sarkar, 2018), whereas NK cells rely on other cells to make the necessary cytokines for proliferation (such as IL-2 and IL-15) and survival (such IL-21) (Fauriat *et al*, 2010). Furthermore, NK cells are more dependent on cytokines such as IL-15 compared to the CD8+ T cells (DeGottardi *et al*, 2016). Therefore, cytokine competition for IL-15 and IL-2 might also explain increased death of mature NK cells in the expansion phase. A test of this hypothesis could be investigation of MCMV-induced NK cell maturation in *Il15+/-* mice.

We quantified the population kinetics of antigen-specific Ly49H+ NK cells during the expansion phase of MCMV infection (<8 days post-infection) using both endogenous and adoptively transferred NK cells. This antigen-specific response was further compared with the kinetics of NK cell maturation in the absence of MCMV infection. Adoptively transferred Ly49H+ NK cells grow at an increased rate compared to the endogenous Ly49H+ NK cells (∼3× faster for Ly6C- cells). A possible explanation of this phenomenon is the stronger competition for resources (e.g. antigens and/or cytokines) that would occur in endogenous NK cells could prevent them from growing as quickly as the less dense population of adoptively transferred NK cells. Interestingly, our estimate of the differentiation rate of the immature CD27+ to mature CD27- NK cells showed negligible effect of MCMV infection on this process. This shows that the signaling processes governing NK cell antigen-specific expansion and cytotoxicity are not impacting transcription and translation of CD27, but rather preferential expansion of certain subsets is creating altered phenotype abundances observed in MCMV infection.

Our stochastic modeling of heterogeneous NK cell clones showed that larger stochastic variations in the mature NK cell populations arising due to higher intrinsic noise fluctuations generated by the interplay between the higher proliferation and cell death processes are important for producing large clones of mature NK cells at 8 days post-infection. This is in contrast with stochastic kinetics of CD8+ T cells where the larger clones of effector CD62L- CD8+ T cells can arise due to a higher rate of proliferation of these cells even in the absence of any cell death (Buchholz *et al*., 2013). Our effort to use different models to quantitatively describe the heterogeneity of NK cell clones demonstrate the critical role of the correlation between the clone size and the proportion of the mature NK cells in the clones, *C*_size-CD27+_, in screening out mechanisms that are unable to give rise to the observed heterogeneity.

## Materials and Methods

### Calculation of the first and second moments and *C_size-CD27+_* for the NK cell subpopulations in the clones

The first and the second moments for the number of immature 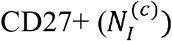 and mature CD27-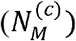 NK cells in *n_clone_* number of clones ({c}) are defined as follows:

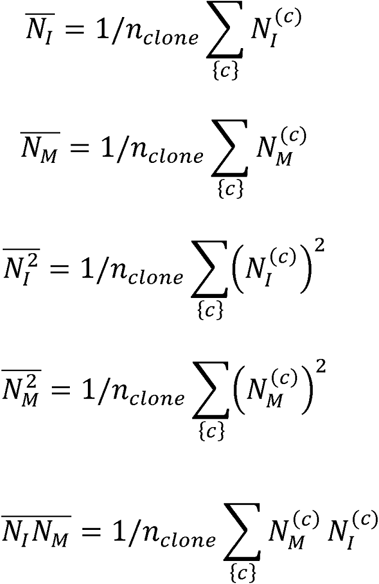

The correlation *C_size-CD27+_* is defined as,

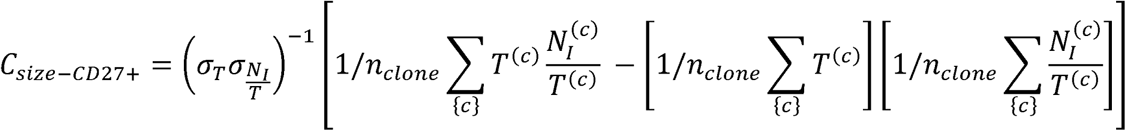

where, 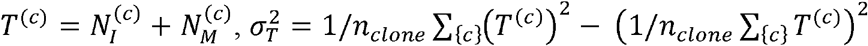 and, 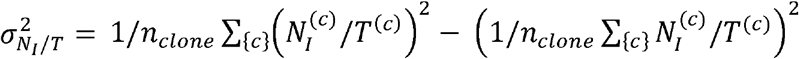.

### NK cell development model

We modeled the NK cell proliferation, death, and maturation as a continuous time Markov processes where a proliferation or a death event in a NK cell population increases or decreases the population number, respectively, by one cell. A differentiation event changing a NK cell from immature (*I*) to mature (*M*) decreases the NK cell population in state *I* and increases the population in state *M* simultaneously by a single NK cell. The kinetics of the system are given by the Master equation describing the time evolution of the conditional probability for the system to be in a particular state containing different numbers of immature and mature NK cells at any time *t* given a specific state at *t=0*. The ODEs describing the kinetics of first and second moments can be derived from the Master equation. The solutions are shown in the Supplementary Materials. The first and second moments for the clone size distributions for all our models were calculated by solving the corresponding ODEs.

### Stochastic simulations

We consider the cell proliferation, differentiation, and death processes as continuous time Markov processes which are modeled as simple birth (n # of cells → n+1 # of cells), death (n# of cells → n-1 # of cells), and differentiation (N_I_ # of immature cells, N_M_ # of mature cells → N_I_ - 1 # of immature cells, N_M_ +1 # of mature cells) processes. The propensities for the above transitions are computed for a range of rates described in Table 1, tables in the supplement, and in the parameter scan section. We use the Gillespie algorithm(Gillespie, 1977) for all our stochastic simulations. For two-stage and three-stage models, we initialized the simulation with a single CD27+ NK cell. We evolved the system until 8 days and numbers of CD27+ and CD27- cells in the NK cell clones were recorded. We calculated the correlation *C*_size-CD27+_ for the clones by averaging over 10^4^ stochastic trajectories.

### Two-state model with differentiation rate increasing with cell division number

This is a variant of a Cyton model where the rates of proliferation, differentiation, and death individual cells are inherited from mother to the daughter cells in dividing cells. In our model, we considered a two-state model where immature CD27+ NK cells differentiate into mature CD27- NK cells, and both immature and mature NK cells replicate with rates k_I_ and k_M_ (k_I_>k_M_). In the model, we tracked the number of divisions experienced by an individual immature cell where the rate of differentiation is proportional to the number of divisions (n_division_) the cell has undergone, i.e., the rate of differentiation in the cell after n_division_ is r_diff_ =r_0_ × n_division_, where r_0_ is the differentiation rate in the founder cell after the first division (Fig. S3). We do not consider any cell death in the model. We implement a continuous time Monte Carlo simulation following Gillespie’s approach to simulate cell proliferation and differentiation in the system. More computational details of this multi-scale method that tracks single-cell divisions can be found in Wethington *et. al* (Wethington *et al*, 2025). We adapted the framework (Wethington *et al*., 2025) to include: 1) a propensity for division of immature cells, 2) a propensity for division of mature cells, and 3) a propensity of differentiation that is linearly related to the number of divisions a cell has undergone.

### Two-state model with log-normally distributed activation times

For the model with log-normally distributed activation times, we assume an NK cell received stimulation at time 8-*t* days and then expanded for a duration of *t* days until day 8. We assign based on the distribution

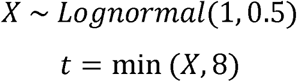

Each clone is evaluated for a randomly sampled *t* according to this distribution.

### Fitting model to moments

The first and second moments 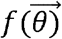 for the clone size distributions obtained for two- and three-stage models were computed by solving ODEs (Eqs. S2-S6 in Supplementary Text 1 and the ODEs in Supplementary Text 2). The parameter vector 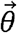 represents the rates for proliferation, differentiation, and death in the models. To compare the predicted moments 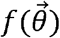 from the model with the actual moments *y* computed from the data, we used the cost function below:

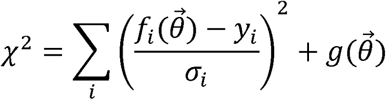

where, σ*_i_* is the standard deviation corresponding to the data used for computing *y_i_*. A penalty function, 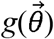 is introduced to implement the constrained imposed by sign of the correlation Csize*_-CD27+_*. We set 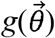 to a high value (= 10^4^) when *C_size-CD27+_* > -0.2 at any value of Δ*_k_* or *C_size-CD27+_* < 0 and Δ*_k_* < 0, otherwise 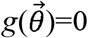. This constraint describes the experimental observation that immature CD27- NK cells grow faster than mature CD27+ NK cells in the expansion phase. σ*_i_* is calculated by taking the standard deviation of each moment across 10,000 bootstrapped samples of the clones. Minimization of the cost function was performed with the *lmfit* python package, using the ‘least_squares’ Trust Region Reflective algorithm. Parameter search space was confined from 0.0 to 2.0 day^-1^ for all parameters. We also performed a parameter fitting without any constraint on Δ_k_, instead fitting the ratio of cells arising from a 1:1 mix of CD27+/CD27- NK cells following the experiments in Flommersfeld *et al.(Flommersfeld et al., 2021)*, adding one additional data point and residual to the model. The resulting best-fit parameters represent Δ_k_ <0, thus we reasoned these estimated parameters do not capture the NK cell expansion observed in experiments (Table S6). This is in line with the previous findings that setting CD27- as the faster growing population can match observed moments data, but this must be balanced with the finding that CD27+ cells grow faster (i.e. Δ_k_>0).

### Effective Growth Rate for 3-stage system

We developed a metric to quantify the difference in the growth rate of immature CD27- and mature CD27+ NK cells for the three-stage model. We computed mean populations of the NK cell subsets at day 8 by solving Eqs. S17-S25 in Supplementary Text 2 for two different initial conditions. In one case we initialized with the immature CD27+Ly6C- cells, and in the other case we initialized with equal proportions of mature CD27-Ly6C- and mature CD27-Ly6C+ cells. We then computed the “effective growth difference” (Δ*_k_*) by evaluating the difference in the mean populations of the immature CD27+ and the mature CD27- NK cells at day 8. We performed an asinh(Δ_k_) transformation to help visualization of the data in the graphs shown in the manuscript.

### Parameter scans

For parameter scans showing the relationship between growth rates and the correlation between clone size and %CD27+ cells, each parameter is randomly selected from a uniform distribution from 0 to 1.6 day^-1^. For each parameter set, 10^4^ Gillespie simulations were carried out to compute the correlation *C*_size-CD27+_ for the NK cell clones at day 8. We performed scans for 10^5^ different parameter sets.

### Ordinary differential equation fits to homeostatic data

The growth and differentiation kinetics of populations of CD27+Tom+ (n_1_), CD27-Tom+ (n_2_), CD27+Tom- (n_3_), and CD27-Tom- (n_4_) cells in the periphery in homeostasis are given by the ODEs below:

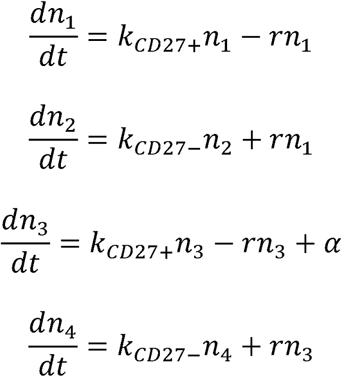

The CD27+ and CD27- cells grow with net rates k_CD27+_ and k_CD27-_, respectively, and the differentiation CD27+ → CD27- occurs at a rate, *r*. The net rates (k_CD27+_ and k_CD27-_) of growth or loss relate to clonal rather than cellular dynamics. The influx of immature CD27- cells from the bone marrow to the periphery occurs at a rate α. At homeostasis it is reasonable to assume that the sum of all Ly49H+ and the sum of all Ly49H- NK cells were assumed to be constant through time, *i.e*.,

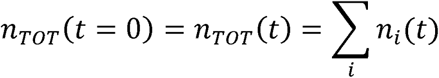

The kinetics of the percentages (N_i_ = 100×(n_i_/n_TOT_)) of each cell type follow the ODEs below:

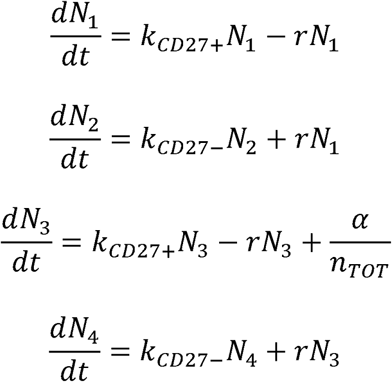

We define 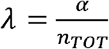, which describes the influx rate of either the total Ly49H+ or total Ly49H-population as a percentage of the given population. This makes the ODEs which we fit:

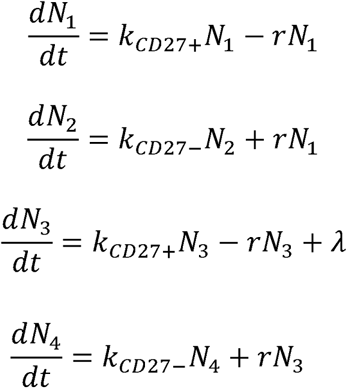

We assume the total number of NK cells and the relative abundances of varying subsets do not change with time in homeostasis, therefore,

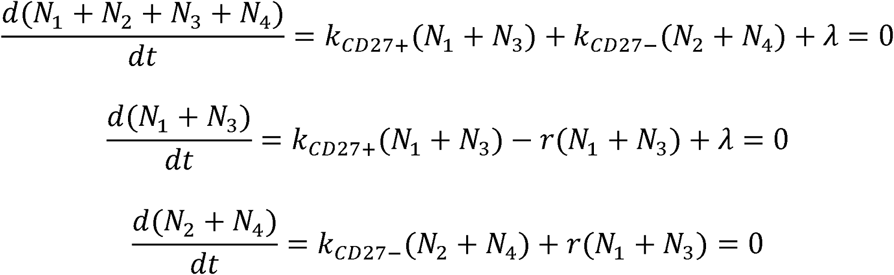

These constraints lead to equalities between parameters,

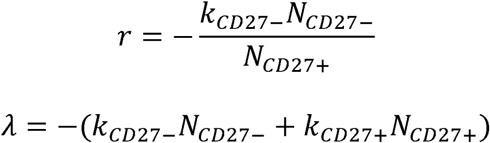

Thus, by estimating only *k_CD27+_* and *k_CD27-_*, and using the mean relative abundances of CD27+ and CD27- cells across mice and time, we can solve all four rate parameters in the homeostatic state.

### Ordinary differential equation fits to mass cytometry data

We gated Ly49H+ NK cells in the CyTOF studies into three subsets: CD27+Ly6C-, CD27- Ly6C-, and CD27-Ly6C+. Details of the gating are described in the Supplementary Material. We describe the net growth and differentiation kinetics of the populations the immature CD27+Ly6C- (N_I_), mature CD27-Ly6C- (N_INT_), and mature CD27-Ly6C+ (N_M_) by the ODEs below,

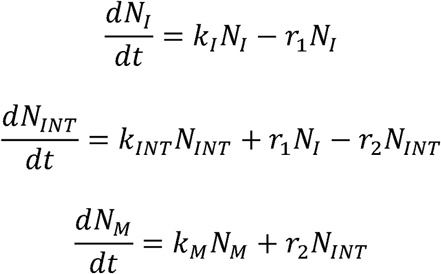

Where, *k_I_*, *k_INT_*, and *k_M_* denote the net rates of growth (or loss) for *N_I_*, *N_INT_*, and *N_M_*, respectively. The net rates (*k_I_*, *k_INT_* and *k_M_*) of growth or loss relate to clonal rather than cellular dynamics. The rates of differentiation for CD27+Ly6C- → CD27-Ly6C- and CD27-Ly6C- → CD27- Ly6C+ are given by *r_1_* and *r_2_*, respectively. Since the above system of ODEs is autonomous and linear, we can obtain the time dependent solutions analytically and exactly. The analytical solution was numerically evaluated for different values of the rate parameters represented by 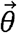. The analytical solutions were evaluated numerically for specific rate constants and initial conditions. We used the estimated population sizes (details in the main text) for the above NK cell subsets in the CyTOF data at t=0 as the initial conditions and computed the population sizes at day 4 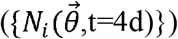 and day 7 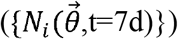 by solving the above ODEs for a parameter set 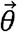. We then estimated the parameters by 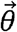 by fitting the population sizes at day 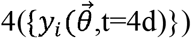 and day 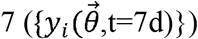 using the cost function below.

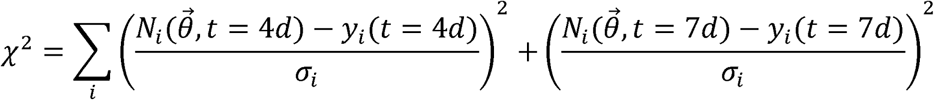

### Bootstrapping confidence intervals for parameter estimates

To assess the confidence intervals for parameter estimates, we randomly sampled the clones with replacement with equal probability, calculated each moment and its standard deviation, and fit the moments as described above. We repeated the process 1,000 times, yielding 1,000 estimates for the parameters θ. We then sorted the parameter estimates by magnitude and took the 2.5^th^ and 97.5^th^ percentile for each. These estimates made up the 95% confidence intervals reported here.

For estimations of the homeostatic and endogenous NK populations, we sampled a population of mice before sampling errors in measurements for each bootstrap estimate.

### Bootstrapping confidence intervals for model fits

The model fits in Figures 3a and 3b have confidence intervals around them. These are calculated by bootstrapping parameter estimates as described above, and simulating a trajectory for each bootstrapped parameter estimate set. At each time *t*, we take the 2.5^th^ and 97.5^th^ percentile of the model trajectory for each cell type. These make up the lower and upper limit for the 95% confidence interval at time *t*.

### Animals and MCMV infection

C57BL/6 (B6) B6 mice were purchased from the National Cancer Institute at 6 weeks of age and housed in the specific pathogen-free animal facility at the University of California San Francisco (UCSF) in accordance with the guidelines of the Institutional Animal Care and Use Committee. Experiments used mice of both genders, aged 8-12 weeks.

B6 mice were infected with 1000 plaque-forming units (PFU) of Smith strain MCMV via intraperitoneal injection. Spleens were acquired different time point following MCMV infection and examined by flow cytometry.

### Flow Cytometry

Splenocytes from uninfected or infected mice were processed into single cell suspensions, RBC lysed, and then stained for flow cytometry. Surface staining was performed using pre-conjugated antibodies in the presence of the viability marker ZombieRed (BioLegend) and were then analyzed on an LSR II cytometer (BD Biosciences). The antibodies used for flow cytometry were: CD3ε (145-2C11), CD19 (6D5), CD49b (DX5), NK1.1 (PK136), KLRG1 (2F1), Ly6C (HK1.4), CD27 (LG.3A10), and Ly49H (3D10). All antibodies were purchased from BioLegend, BD Biosciences, or ThermoFisher Scientific. An unconjugated anti-CD16/CD32 antibody (clone 2.4G2, UCSF Antibody Core) was used to block non-specific binding.

### Mass cytometry

Samples were prepared for mass cytometry as previously described (Potempa *et al*., 2022). Briefly, splenocytes from each mouse were obtained by mechanical dissociation through a 40-μm filter, treated with ACK lysis buffer, and resuspended in PBS + 5 mM EDTA (PBS/EDTA). Cells were stained for viability by mixing 1:1 with PBS/EDTA + 100 mM cisplatin (Enzo Life Sciences) for 1 minute before quenching with PBS/EDTA + 0.5% BSA (Staining Media). One million splenocytes from each mouse was stained with metal-conjugated antibodies against CD16 (Nd144) and CD32 (Eu153) for 15 minutes at room temperature in Staining Media before supplementation with an Fc blocking Ab (clone 2.4G2). After washing in Staining Media, cells were stained with a master mix of 35 additional metal-conjugated antibodies (Table S7) for 30 minutes at room temperature. At the conclusion of the staining period, cells were washed in PBS/EDTA before fixation with PBS/EDTA + 3.2% paraformaldehyde and stored at 4° C for 3-5 days. One day prior to data acquisition, samples were barcoded using a 20-plex Pd barcoding kit (Fluidigm) per manufacturer protocol, pooled, and fixed again, overnight at 4° C, in a PBS/EDTA + 3.2% paraformaldehyde solution supplemented with a 1:1000 dilution of a 191/193 DNA intercalator (Fluidigm). Just prior to acquisition, barcoded samples were pelleted, washed in double-deionized water, and resuspended in double-deionized water with a 1:20 dilution of EQ four Element Calibration Beads (Fluidigm). Data were acquired using a CyTOF2- Helios mass cytometer (Fluidigm), which were then normalized and debarcoded prior to analysis.

## Acknowledgements

This work was supported by NIH grant R01AI146581 to J.D. N.M.A. was supported by a Medical Scientist Training Program grant from the National Institute of General Medical Sciences (T32GM007739 to the Weill Cornell/Rockefeller/Sloan Kettering Tri-Institutional MD-PhD Program) and by an F30 Predoctoral Fellowship from the National Institute of Allergy and Infectious Diseases (F30 AI136239). We would like to thank the Ohio Supercomputer Center for computational resources.

