## Supplementary Materials for "Clonal stochasticity in early NK cell response to mouse cytomegalovirus is generated by mature subsets of varying proliferative ability"

**Supplementary Text 1: Analysis of the stochastic kinetics of NK clonal expansion**

Two-Stage System

We consider the model shown in Fig. 1a. The immature (state I) CD27+ NK cells differentiate into mature (state M) CD27- NK cells with a rate *r*, i.e., $I\underset{\to}{r}M$ . The immature and mature NK cells proliferate with rates $k_{I}$ and $k_{M}$, described by the reactions $I\underset{\to}{b_{I}}I+I$ and$M\underset{\to}{b_{M}}M+M$ , respectively. We will also consider a model with cell death described by reactions $I\underset{\to}{d_{I}}\phi$ and$M\underset{\to}{d_{M}}\phi$. The stochastic kinetics of these processes are given by the Master equation describing the time evolution of the probability distribution $P_{N_{I},N_{M}}\left( t \right)$of the numbers of immature (N_I_) and mature (N_M­_) NK cells at any time *t* given the initial condition $N_{I}\left( t=0 \right)=1, N_{M}\left( t=0 \right)=0$. Note that the model without death may be described as the model with death with $d_{I}=d_{M}=0$.

We can easily derive the master equation for $P_{N_{I},N_{M}}(t)$:

$\frac{dP_{N_{I},N_{M}}}{dt}=b_{I}\left( N_{I}-1 \right)P_{N_{I}-1,N_{M}}+d_{I}\left( N_{I}+1 \right)P\left( N_{I}+1,N_{M} \right)+r\left( N_{I}+1 \right)P_{N_{I}+1,N_{M}-1}+b_{M}\left( N_{M}-1 \right)P_{N_{I},N_{M}-1}+d_{M}\left( N_{M}+1 \right)P_{N_{I},N_{M}+1}-\left( \left( b_{I}+d_{I}+r \right)N_{I}+(b_{M}+d_{M})N_{M} \right)P_{N_{I},N_{M}}$

(S1)

The ODEs describing the average values (indicated by overbars) of N_I_ and N_M_ can be derived from the above Master equation in Eq. (S1).

$$\frac{d\bar{N}_{I}}{dt}=(b_{I}-d_{I}-r)\bar{N}_{I}$$

(S2)

$$\frac{d\bar{N}_{M}}{dt}=r\bar{N}_{I}+(b_{M}-d_{M})\bar{N}_{M}$$

(S3)

2^nd^ order:

$$\frac{d\bar{N_{I}^{2}}}{dt}=2\left( b_{I}-d_{I}-r \right)\bar{N_{I}^{2}}+\left( b_{I}+d_{I}+r \right)\bar{N}_{I}$$

(S4)

$$\frac{d\bar{N_{M}^{2}}}{dt}=2{(b}_{M}-d_{M})\bar{N_{M}^{2}}+{(b}_{M}+d_{M})\bar{N}_{M}+2r\bar{N_{I}N_{M}}+r\bar{N}_{I}$$

(S5)

$$\frac{d\bar{N_{I}N_{M}}}{dt}=\left( b_{I}+b_{M}-d_{I}-d_{M}-r \right)\bar{N_{I}N_{M}}+r\bar{N_{I}^{2}}-r\bar{N}_{I}$$

(S6)

Analysis of the correlation *C*_size-CD27+_

The correlation *C*_size-CD27+_ describes the correlation between the size or the total population ($N_{I}+N_{M}$) of the NK cell clones and the fraction ($\frac{N_{I}}{N_{I}+N_{M}}$) of the immature CD27+ NK cells at any time *t*. For simplicity of notation, we rename *C*_size-CD27+_ as $Corr\left( N_{I}+N_{M},\frac{N_{I}}{N_{I}+N_{M}} \right)$ for this section. The standard deviation *σ_x_* for a variable *x* is given by $\sigma_{x}=\sqrt{\bar{x^{2}}-\bar{x}^{2}}$ ; the overbar represents average over the probability distribution, $P_{N_{I},N_{M}}.$

$$Corr\left( N_{I}+N_{M},\frac{N_{I}}{N_{I}+N_{M}} \right)=\frac{\bar{\left( \left( N_{I}+N_{M}-\bar{\left( N_{I}+N_{M} \right)} \right)\left( \frac{N_{I}}{N_{I}+N_{M}}-\bar{\frac{N_{I}}{N_{I}+N_{M}}} \right) \right)}}{\sigma_{N_{I}+N_{M}}\sigma_{\frac{N_{I}}{N_{I}+N_{M}}}}$$

(S7)

We now evaluate conditions that can make the above correlation negative at t=8 days. Since the denominator in Eq. (S7) is positive, we consider the numerator for the analysis. The numerator can be expanded in terms of fluctuations around the mean values of N_I_ and N­_M_, i.e., $N_{I}(t)=\bar{N_{I}}(t)+\delta N_{I}(t)$ and $N_{M}(t)=\bar{N}_{M}(t)+\delta N_{M}(t)$, as follows

$$=\bar{N}_{I}-\left( \bar{N}_{I}+\bar{N}_{M} \right)\bar{\left( \frac{\frac{\bar{N}_{I}+\delta N_{I}}{\bar{N}_{I}+\bar{N}_{M}}}{1+\frac{\delta N_{I}+\delta N_{M}}{\bar{N}_{I}+\bar{N}_{M}}} \right)}$$

(S8)

Now, considering fluctuations where $\frac{\delta N_{I}+\delta N_{M}}{\bar{N}_{I}+\bar{N}_{M}}<1$, we expand the above expression to the second order as,

$$\approx\bar{N}_{I}-\left( \bar{N}_{I}+\bar{N}_{M} \right)\bar{\left( \left( \frac{\bar{N}_{I}+\delta N_{I}}{\bar{N}_{I}+\bar{N}_{M}} \right)\left( 1-\frac{\delta N_{I}+\delta N_{M}}{\bar{N}_{I}+\bar{N}_{M}}+\left( \frac{\delta N_{I}+\delta N_{M}}{\bar{N}_{I}+\bar{N}_{M}} \right)^{2} \right) \right)}$$

(S9)

Reducing and noting that $\bar{\delta N_{I}}$ and $\bar{\delta N_{M}}$ are zero,

$$=\frac{1}{\left( \bar{N}_{I}+\bar{N}_{M} \right)^{2}}\frac{\bar{N}_{M}\delta\bar{N_{I}^{2}}+\left( \bar{N}_{M}-\bar{N}_{I} \right)\bar{\delta N_{I}\delta N_{M}}-\bar{N}_{I}\delta\bar{N_{M}^{2}}}{\sigma_{\frac{N_{I}}{N_{I}+N_{M}}}\sigma_{N_{I}+N_{M}}}$$

(S10)

Inputting this in Eq. (S10),

$$Corr(N_{I}+N_{M},\frac{N_{I}}{N_{I}+N_{M}})\approx\frac{1}{\left( \bar{N}_{I}+\bar{N}_{M} \right)^{2}}\frac{\bar{N}_{M}\sigma_{N_{I}}^{2}+\left( \bar{N}_{M}-\bar{N}_{I} \right)Cov(N_{I},N_{M})-\bar{N}_{I}\sigma_{N_{M}}^{2}}{\sigma_{\frac{N_{I}}{N_{I}+N_{M}}}\sigma_{N_{I}+N_{M}}}$$

(S11)

Where

$$Cov\left( N_{I},N_{M} \right)=\bar{N_{I}N_{M}}-\bar{N}_{I}\bar{N}_{M}$$

(S12)

Note$,$ $\bar{N}_{M}-\bar{N}_{I}>0$for the NK cell populations at day 8. We checked the validity of the approximation in Eq. (S11) with our numerical evaluation of the correlation function in Eq. (S7) (Fig. S1). We find that the sign of approximate correlation function in Eq. (S11) is in agreement with the sign of the actual correlation function in Eq. (S7). Therefore, use the approximate form in Eq. (S11) for further analysis. The approximate form of the correlation function in Eq. (S11) can become negative if $\sigma_{N_{M}}^{2}$ is larger than $\sigma_{N_{I}}^{2}+\left( \bar{N}_{M}-\bar{N}_{I} \right)Cov(N_{I},N_{M})$. Below we show that increasing the death rate *d_M_* increases $\sigma_{N_{M}}^{2}$ when the growth rate of the mature population ${k_{M}=b}_{M}-d_{M}$ and the other rates, *b_I_*, *d_I_*, and *r* are held fixed. Eq. (S5) can be recast in terms of *k*_M_ and the other rate parameters as,

$$\frac{d\bar{N_{M}^{2}}}{dt}=2k_{M}\bar{N_{M}^{2}}+(k_{M}+{2d}_{M})\bar{N}_{M}+2r\bar{N_{I}N_{M}}+r\bar{N}_{I}$$

(S13)

Since the variables $\bar{N}_{I}$ and $\bar{N}_{M}$ do not depend on $d_{M}$ when Eqs. S2-S3 are recast in terms of k_M_, $\frac{\partial\bar{N_{M}^{2}}}{\partial d_{M}}$ follows the equation below.

$$\frac{d}{dt}\frac{\partial\bar{N_{M}^{2}}}{\partial d_{M}}=2k_{M}\frac{\partial\bar{N_{M}^{2}}}{\partial d_{M}}+2\bar{N}_{M}$$

(S14)

where the solution is given by

$$\frac{\partial\bar{N_{M}^{2}}}{\partial d_{M}}=\frac{\partial\bar{N_{M}^{2}}}{\partial d_{M}}\left( t=0 \right)+2e^{2k_{M}t}\int_{0}^{t} dt^{'}e^{-2k_{M}t^{'}}\bar{N}_{M}$$

(S15)

We can simplify the solution further as

$\frac{\partial\bar{N_{M}^{2}}}{\partial d_{M}}=\frac{2r}{k_{M}\left( g-k_{M} \right)\left( g-2k_{M} \right)}(k_{M}e^{rt}+\left( g-2k_{M} \right)e^{k_{M}t}-\left( g-k_{M} \right)e^{2k_{M}t})$

(S16)

where,

$$g=b_{I}-d_{I}-r$$

Eq. (S16) shows $\frac{\partial\bar{N_{M}^{2}}}{\partial d_{M}}\left( t=0 \right)=0$, therefore from Eq. (2) $\frac{\partial\bar{N_{M}^{2}}}{\partial d_{M}}\left( t \right)=2\int_{0}^{t} dt^{'}e^{-2k_{M}{(t}^{'}-t)}\bar{N}_{M}(t')\geq0$ as the integrand is always non-negative. Since, for the parameters $\bar{N}_{M}$ would be non-zero for $t\leq8$ days, we can conclude that increasing the death rate $d_{M}$­ would increase the variance $\sigma_{N_{M}}^{2}$ causing the correlation to become negative.

**Numerical Integration of Master equation and calculation of AIC_c_**

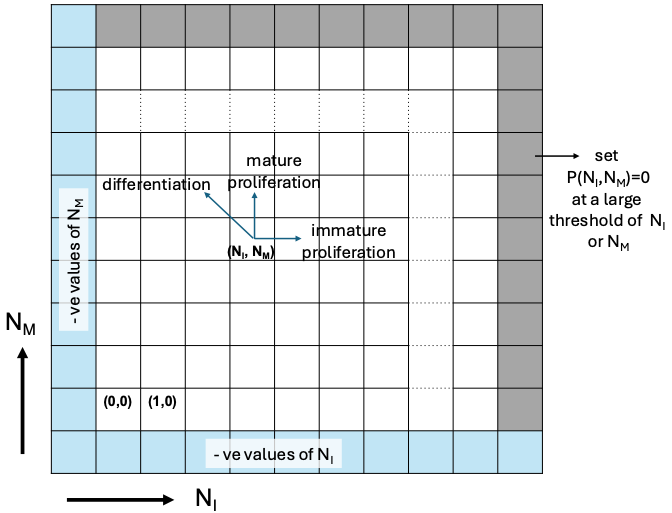
We numerically solve the Master equation in Eq. (S1) which describes the probability distribution $P_{N_{I},N_{M}}\left( t \right)$of the numbers of immature ($N_{I}$) and mature ($N_{M}$) NK cells at any time $t$ given the initial condition $P_{1,0}=1$ at time $t=0$. We use numerical integration method to solve the coupled ordinary differential equations (ODEs) in Eq. (S1). Since there is no upper bound to the number of $N_{I}$ and $N_{M}$, an exact solution of Eq. (S1) would need to consider the ranges $0\leq N_{I}\leq\infty$ and $0\leq N_{M}\leq\infty.$ However, accounting for the above ranges in numerical solution of the ODEs will be computationally infeasible, therefore, we approximate the above system of ODEs by considering large but finite upper bounds of $N_{I}$ and $N_{M}$, i.e., $0\leq N_{I}\leq N_{I}^{(bound)}$ and $0\leq N_{M}\leq N_{M}^{(bound)}$where $N_{I}^{(bound)}$=$N_{M}^{(bound)}=1200$. The transition rates carrying the system from inside the boundary to outside the boundary or vice versa are set to zero. This system will provide a good approximation for the full solution for a time interval when the exact solution of the kinetics has a very low probability to transition across the boundary. We checked this with the exact solution for the pure birth model for the time interval of 0 to 8 days where largest difference in the probability distributions between the exact and the approximate solutions was $<{10}^{-9}($Fig. S1c). We used ‘*odeint’* function with 8^th^ -order Runge-Kutta method ‘*dopri8*’ from ‘*torchdiffeq’* and implemented those in GPUs to numerically solve the ${10}^{6}$ coupled ODEs in Eq. (S1) with the above bounds in N_I_ and N_M_. We then computed the likelihood $L$of the observed $38$ clone sizes (${\{N}_{I}^{(i)}, N_{M}^{(i)}, i=1,..,38\}$) (Fig. S1b) given the numerical solution of $P(N_{I},N_{M};\theta)$ where $\theta$ denotes the parameters $b_{I}, b_{M}, d_{I},d_{M}$ and, $r$. The Likelihood $L(\theta)$ is given by,

Schematic diagram showing the values of the numbers of immature ($N_{I}$) and mature ($N_{M})$ for which the Master equation was solved numerically. The boundary layers ($N_{I}^{ub},{0\leq N}_{M}\leq N_{M}^{ub})$ and $(0{\leq N}_{I}\leq N_{I}^{ub},N_{M}^{ub}$) at the upper bounds of the numbers used to make the computation feasible are marked in grey. The values where $N_{I}$ and $N_{M}$ assume negative values are shown in blue.

$$L(\theta)=\prod_{i=1}^{38} P\left( N_{I}^{\left( i \right)},N_{M}^{\left( i \right)};\theta\right)$$

We find optimal parameters ($\hat{\theta}$) by minimizing $-log(L(\theta)$) (Fig. S1d) using the ‘*minimize’* function with *‘Nelder-Mead’* algorithm from ‘*scipy’.* Then we compute $AICc$ for comparing the models, where $AICc$ for a model with $K$ parameters describing $n$ number of data points is defined as

$$AIC_{c}=AIC+\frac{2K(K+1)}{n-K-1}$$

and the Akaike Information Criterion ($AIC$) is given by $AIC=2K-2log(L(\hat{\theta}))$.

The $log(L(\hat{\theta}))$ values for two state model without death and with death are $-545.4$ and $-514.6,$ , respectively. The corresponding $AICc$values are $1097.5$ and $\text{1041.0}$, respectively. Thus, the model with death has a smaller $AICc$compared to the model without death with a difference of $\Delta AICc >50,$ clearly demonstrating that the model with death is favored over the other model by the observed data.

**Supplementary Text 2: Moment equations for the three state and other models**

Three-Stage Model

$$\frac{d\bar{N}_{I}}{dt}=(b_{I}-d_{I}-r_{1})\bar{N}_{I}$$

(S17)

$$\frac{d\bar{N}_{INT}}{dt}=r_{1}\bar{N}_{I}+(b_{INT}-d_{INT}-r_{2})\bar{N}_{INT}$$

(S18)

$$\frac{d\bar{N}_{M}}{dt}=r_{2}\bar{N}_{INT}+(b_{M}-d_{M})\bar{N}_{M}$$

(S19)

$$\frac{d\bar{N_{I}^{2}}}{dt}=2\left( b_{I}-d_{I}-r_{1} \right)\bar{N_{I}^{2}}+(b_{I}+d_{I}+r_{1})\bar{N}_{I}$$

(S20)

$$\frac{d\bar{N_{INT}^{2}}}{dt}=2\left( b_{INT}-d_{INT}-r_{2} \right)\bar{N_{INT}^{2}}+{(b}_{INT}+d_{INT}+r_{2})\bar{N}_{INT}+2r_{1}\bar{N_{I}N_{INT}}+r_{1}\bar{N}_{I}$$

(S21)

$$\frac{d\bar{N_{M}^{2}}}{dt}=2\left( b_{M}-d_{M} \right)\bar{N_{M}^{2}}+{(b}_{M}+d_{M})\bar{N}_{M}+2r_{2}\bar{N_{INT}N_{M}}+r_{2}\bar{N}_{INT}$$

(S22)

$$\frac{d\bar{N_{I}N_{INT}}}{dt}=\left( b_{I}+b_{INT}-d_{I}-d_{INT}-r_{1}-r_{2} \right)\bar{N_{I}N_{INT}}+r_{1}\bar{N_{I}^{2}}-r_{1}\bar{N}_{I}$$

(S23)

$$\frac{d\bar{N_{INT}N_{M}}}{dt}=\left( b_{INT}+b_{M}-d_{INT}-d_{M}-r_{2} \right)\bar{N_{INT}N_{M}}+r_{2}\bar{N_{INT}^{2}}-r_{2}\bar{N}_{INT}+r_{1}\bar{N_{I}N_{M}}$$

(S24)

$$\frac{d\bar{N_{I}N_{M}}}{dt}=\left( b_{I}+b_{M}-d_{I}-d_{M}-r_{1} \right)\bar{N_{I}N_{M}}+r_{2}\bar{N_{I}N_{INT}}$$

(S25)

To change the above system to the model with 3-stages but the intermediate stage is CD27+, simply replace the parameters *b_INT_* and *d_INT_* with *b_DP_* and *d_DP_*.

Two-stage model with asymmetric division

$$\frac{d\bar{N}_{I}}{dt}=(b_{sym}-d_{I}-r)\bar{N}_{I}$$

(S26)

$$\frac{d\bar{N}_{M}}{dt}=(r+b_{asym})\bar{N}_{I}+(b_{M}-d_{M})\bar{N}_{M}$$

(S27)

$$\frac{d\bar{N_{I}^{2}}}{dt}=2\left( b_{sym}-d_{I}-r \right)\bar{N_{I}^{2}}+(b_{sym}+d_{I}+r)\bar{N}_{I}$$

(S28)

$$\frac{d\bar{N_{M}^{2}}}{dt}=2\left( b_{M}-d_{M} \right)\bar{N_{M}^{2}}+{(b}_{M}+d_{M})\bar{N}_{M}+2(r+b_{asym})\bar{N_{I}N_{M}}+(r+b_{asym})\bar{N}_{I}$$

(S29)

$$\frac{d\bar{N_{I}N_{M}}}{dt}=\left( b_{sym}+b_{M}-d_{I}-d_{M}-r \right)\bar{N_{I}N_{M}}+(r+b_{asym})\bar{N_{I}^{2}}-r\bar{N}_{I}$$

(S30)

Two-stage model with inflow

$$\frac{d\bar{N}_{DP}}{dt}=a+(b_{DP}-d_{DP}-r)\bar{N}_{DP}$$

(S31)

$$\frac{d\bar{N}_{M}}{dt}=r\bar{N}_{DP}+(b_{M}-d_{M})\bar{N}_{M}$$

(S32)

$$\frac{d\bar{N_{DP}^{2}}}{dt}=2\left( b_{DP}-d_{DP}-r \right)\bar{N_{DP}^{2}}+{(b}_{INT}+d_{INT}+r+2a)\bar{N}_{DP}+a$$

(S33)

$$\frac{d\bar{N_{M}^{2}}}{dt}=2\left( b_{M}-d_{M} \right)\bar{N_{M}^{2}}+{(b}_{M}+d_{M})\bar{N}_{M}+2r\bar{N_{DP}N_{M}}+r\bar{N}_{DP}$$

(S34)

$$\frac{d\bar{N_{DP}N_{M}}}{dt}=\left( b_{INT}+b_{M}-d_{INT}-d_{M}-r \right)\bar{N_{DP}N_{M}}+r\bar{N_{DP}^{2}}-r\bar{N}_{DP}+a\bar{N}_{M}$$

(S35)

**Table S1:** Best fit parameters to 2-stage model with birth and death describing NK cell clonal bursts given in Fig. 1e.

| **Parameter** | **Estimate** |
| --- | --- |
| b_I_ | 0.662 day^-1^ |
| b_M_ | 1.441 day^-1^ |
| d_I_ | 0.129 day^-1^ |
| d_M_ | 0.964 day^-1^ |
| r | 0.135 day^-1^ |

**Table S2:** Best fit parameters to asymmetric division model describing NK cell clonal bursts given in Fig. S5a.

| **Parameter** | **Estimate**  **(confidence interval)** |
| --- | --- |
| b_sym_ | 0.716 day^-1^  (0.704 – 0.811) |
| b_asym_ | 0.708 day^-1^  (0.663 – 0.725) |
| b_M_ | 1.406 day^-1^  (1.362 – 1.442) |
| d_I_ | 0.039 day^-1^  (0.026 - 0.097) |
| d_M_ | 0.432 day^-1^  (0.401 – 0.481) |
| r | 0.017 day^-1^  (0.004 - 0.025) |

**Table S3:** Best fit parameters to 3-stage model with two CD27+ stages describing NK cell clonal bursts given in Fig. S4a.

| **Parameter** | **Estimate**  **(confidence interval)** |
| --- | --- |
| b_I_ | 0.899 day^-1^  (0.803 – 0.963) |
| b_DP_ | 1.450 day^-1^  (1.247 - 1.801) |
| b_M_ | 0.995 day^-1^  (0.992 – 2.000) |
| d_I_ | 0.000 day^-1^  (0.000 - 0.007) |
| d_DP_ | 0.004 day^-1^  (0.000 - 0.271) |
| d_M_ | 0.076 day^-1^  (0.067 – 1.055) |
| r_1_ | 0.017 day^-1^  (0.000 - 0.018) |
| r_2_ | 0.591 day^-1^  (0.418 - 0.962) |

**Table S4:** Best fit parameters to 2-stage model with constant inflow describing NK cell clonal bursts given in Fig. S3a.

| **Parameter** | **Estimate** |
| --- | --- |
| b_I_ | 1.464 day^-1^ |
| b_M_ | 1.596 day^-1^ |
| d_I_ | 0.332 day^-1^ |
| d_M_ | 0.521 day^-1^ |
| r | 0.263 day^-1^ |
| a | 0.886 cell/day |

**Table S5:** Best fit parameters to endogenous NK cells responding to MCMV infection.

| **Parameter** | **Estimate**  **(confidence interval)** |
| --- | --- |
| k_I_ | 0.108 day^-1^  (0.034 - 0.239) |
| k_INT_ | 1.775 day^-1^  (0.018 – 2.000) |
| k_M_ | -2.000 day^-1^  (-2.000 - 0.367) |
| r_1_ | 3.85×10^-18^ day^-1^  (0.000 – 0.106) |
| r_2_ | 1.482 day^-1^  (0.000 – 1.720) |

**Table S6:** Best fit parameters to 3-stage model with two CD27- stages describing NK cell clonal bursts given in Fig. 2a, but while fitting to comparative growth rates of CD27+ vs. CD27- cells rather than constraining to Δ_k_ > 0. Because r_2_ ~= 0 and Δ_k_ < 0, we report the constrained fit in the main text.

| **Parameter** | **Estimate** |
| --- | --- |
| b_I_ | 1.375 day^-1^ |
| b_INT_ | 2.000 day^-1^ |
| b_M_ | 0.263 day^-1^ |
| d_I_ | 0.350 day^-1^ |
| d_INT_ | 0.866 day^-1^ |
| d_M_ | 2.000 day^-1^ |
| r_1_ | 0.015 day^-1^ |
| r_2_ | 0.000 day^-1^ |

**Table S7**: Mass cytometry antibody panel

| **Metal** | **Channel** | **Target** |
| --- | --- | --- |
| In | 113 | CD45 |
| In | 115 | --- |
| La | 139 | CD62L |
| Pr | 141 | Ly6C |
| Nd | 142 | CD132 |
| Nd | 143 | NKR-P1B |
| Nd | 144 | CD16 |
| Nd | 145 | --- |
| Nd | 146 | CD96 |
| Sm | 147 | 2B4 |
| Nd | 148 | IFNAR |
| Sm | 149 | DNAM-1 |
| Nd | 150 | CD127 |
| Eu | 151 | Ly49D |
| Sm | 152 | CD49b |
| Eu | 153 | CD32 |
| Sm | 154 | CD69 |
| Gd | 155 | CD8 |
| Gd | 156 | CEACAM-1 |
| Gd | 157 | CD3 |
| Gd | 158 | NKG2ACE |
| Tb | 159 | Ly49H |
| Gd | 160 | NK1.1 |
| Dy | 161 | TIM-3 |
| Dy | 162 | --- |
| Dy | 163 | TIGIT |
| Dy | 164 | CD2 |
| Ho | 165 | CD11b |
| Er | 166 | CD137 |
| Er | 167 | CD26 |
| Er | 168 | Ly49CI |
| Tm | 169 | NKp46 |
| Er | 170 | NKG2D |
| Yb | 171 | LAG-3 |
| Yb | 172 | KLRG1 |
| Yb | 173 | CD19 |
| Yb | 174 | Ly49G2 |
| Lu | 175 | IL18Ra |
| Yb | 176 | CD90 |
| Bi | 209 | CD27 |

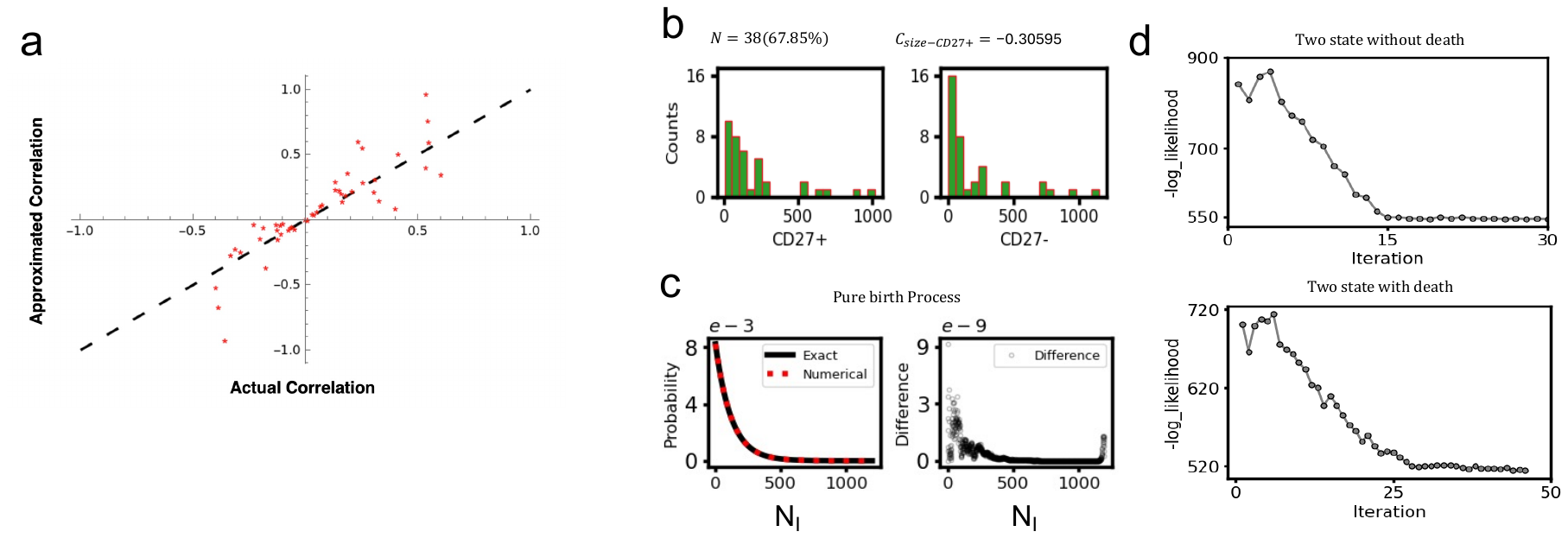

**Figure S1: Analysis of the stochastic kinetics using the Master equation. (a)** *The approximation for the correlation C_size-CD27+_ maintains the sign of the actual correlation.* The approximation for *C_size-CD27+_* (y-axis) was compared with the empirical solution (x-axis) for several parameter sets. Each parameter set is represented by a red star. The black dashed line represents the line *y=x*. For each parameter set, empirical *C_size-CD27+_* was calculated by simulating 1000 clones. **(b)** Histogram of counts of CD27+ and CD27- NK cells shown for the 38 data points used in estimating the model parameters. **(c)** (left) Comparison of the numerical solution of the Master equation at t=8 days with upper bounds in N_I_ for a pure birth process (N_I_ → N_I_+1, rate b_I_) (dashed red) with the exact solution (solid black) with no upper bound in N_I_. The system starts at t=0 with N_I_=1; b_I_ is taken as the largest value taken in our parameter range. (right) Shows the absolute difference in the probability distribution function of the exact solution and numerical solution. The largest difference is of the order of ${10}^{-9}$. **(d)** Shows changes in the -log-likelihood with parameter iterations for the model without death (top) or with death (bottom). The minimum log-likelihood values correspond to -545.4 (top) and -514.6 (bottom). The optimal parameter values are $b_{I}=0.97, b_{M}=1.07, r=0.05$ for two state without death, and, $b_{I}=0.97, b_{M}=1.7, d_{I}=0.15, d_{M}=0.92$ and $r=0.05$ for two states with death.

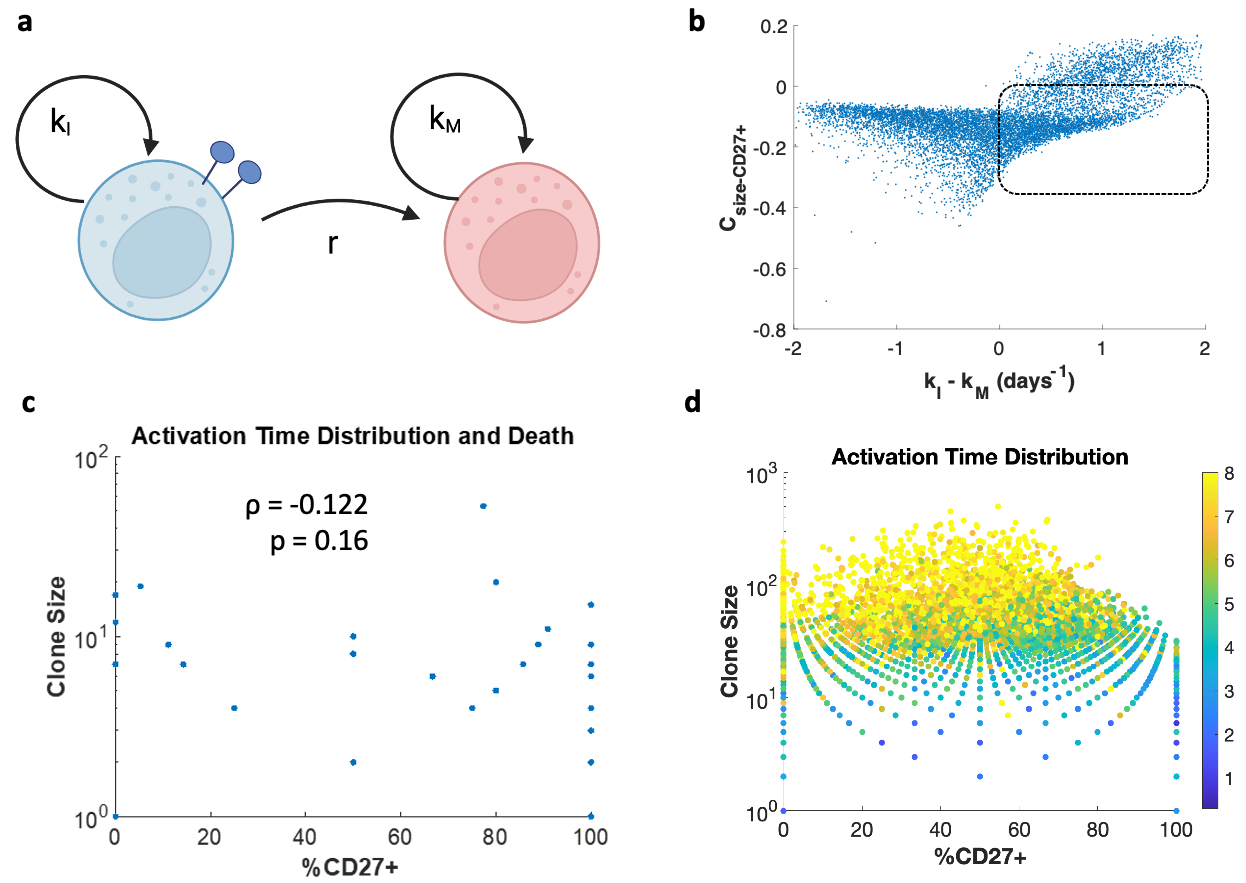

**Figure S2:** **Stochasticity in total activation time helps contribute to negative correlations**. **(a)** The model in (a) differs from the original 2-stage model investigated in that new CD27+ cells are introduced at varying times during the course of infection. We hypothesized that a model where each clone was stimulated for a random length of time drawn from a lognormal distribution could recreate negative correlations as well. Thus, we propose this 2-stage model representation, which is identical to that in Fig. 1a, but with each clone being simulated for a lognormal random length of time. **(b)** Parameter scan for the model in (a). **(c)** Representation of clonal burst simulations for a parameter configuration which generates positive Csize-CD27+. Parameters are taken from Table S1 but with growth rates in place of separate birth and death processes, where kI = 0.533, kM = 0.477, and r = 0.135. The growth rate of CD27+ cells is higher than that of CD27- cells, yet the correlation is -0.122. **(d)** More clonal bursts for the same parameter set as in (c), to demonstrate how stochasticity in total activation time contributes to negative correlations. Color represents the length of activation, so clones that underwent division and differentiation for only one day are shown in blue.

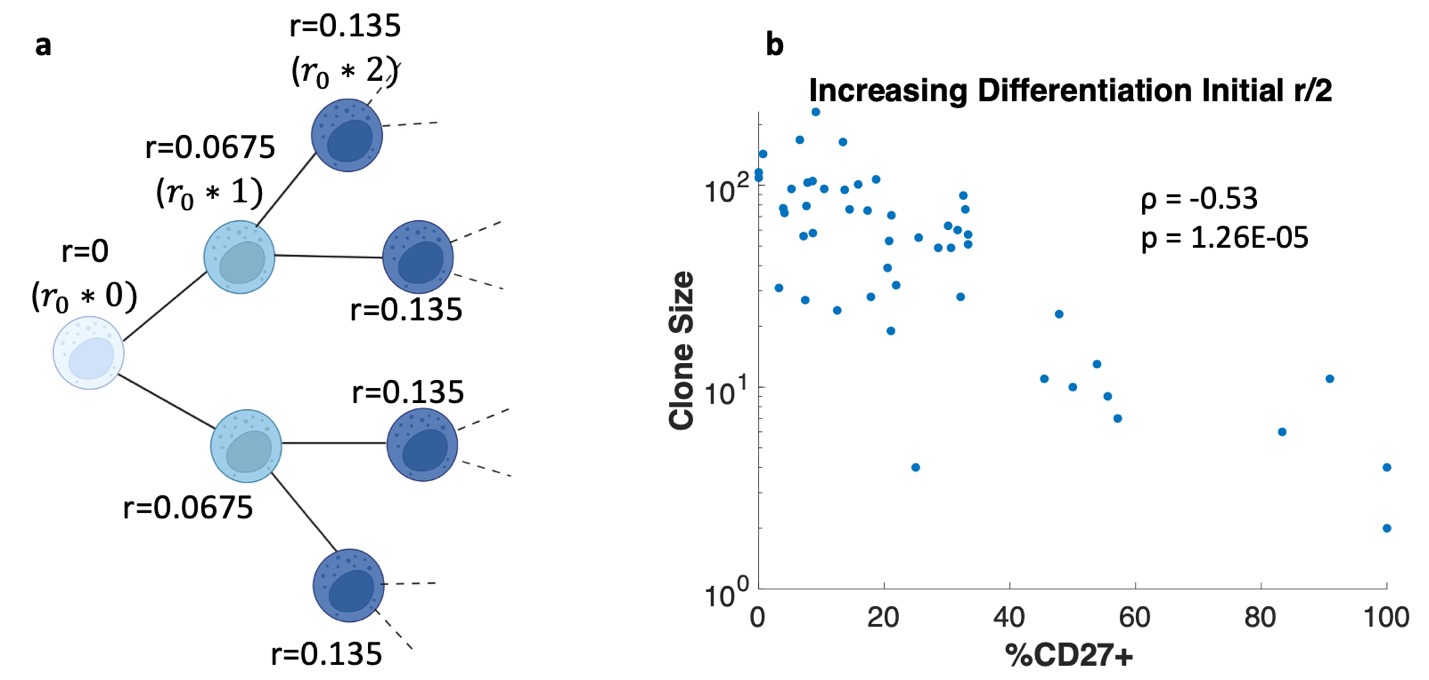

**Figure S3: Linearly increasing differentiation rate with number of divisions results in negative correlations for a 2-stage model. (a)** A schematic representation of our model. The clonal population is initiated by a single founder cell. The daughter cells differ from the mother cell but inherit some attributes. In our case, at each cell division, the rate of differentiation in the daughter cells increases linearly increases from that of the mother cell. In this representation, the rate of differentiation after the first division, r_0_=0.0675, and the rate of differentiation in the daughter cells after n_divisions_ divisions is given by, r_cell_=r_0_×n_divisions_. **(b)** Variation of the clone sizes with the % of CD27+ cells in the clones for 56 clones generated in our simulation of the model in shown (a). Parameters are taken from Table S1 but with growth rates in place of separate birth and death processes, where k_I_ = 0.533, k_M_ = 0.477, and r_0_ = 0.0675. The growth rate of CD27+ cells is higher than that of CD27- cells, yet the correlation is -0.532.

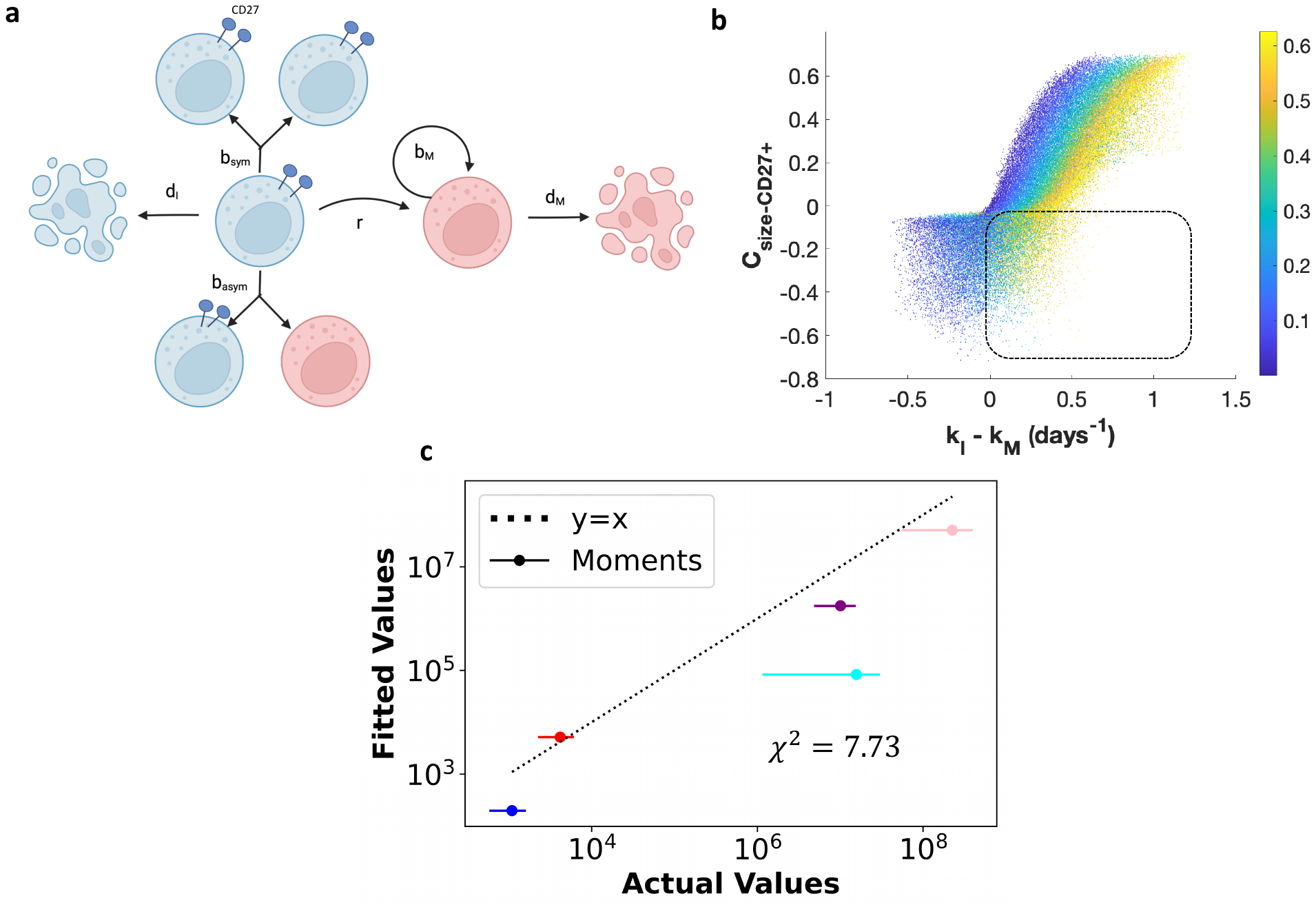

**Figure S4: Modeling asymmetric division of CD27+ cells in a two-stage model.** **(a)** Schematic representation of two-stage model with asymmetric division. It is altered from the model in Fig. 1f so that it has two different terms for division of immature cells. Immature (CD27+) cells can divide symmetrically as before according to rate *b_sym_*, and additionally can divide asymmetrically into a single immature and single mature (CD27-) cell according to rate *b_asym_*. In this model, the growth rates are defined as *k_I_=b_sym_+b_asym_-d_I_* and *k_M_=b_M_-d_M_*. **(b)** A parameter scan for this model where *d_I_=d_M_=0* shows that adding asymmetric division can also account for negative correlations. Each spot represents a unique parameter configuration, with color indicating the value of *b_asym_*. **(c)** Best fit of this model to the moments with constraints Δ_k_ > 0 and C_size-CD27+_ < -0.2.

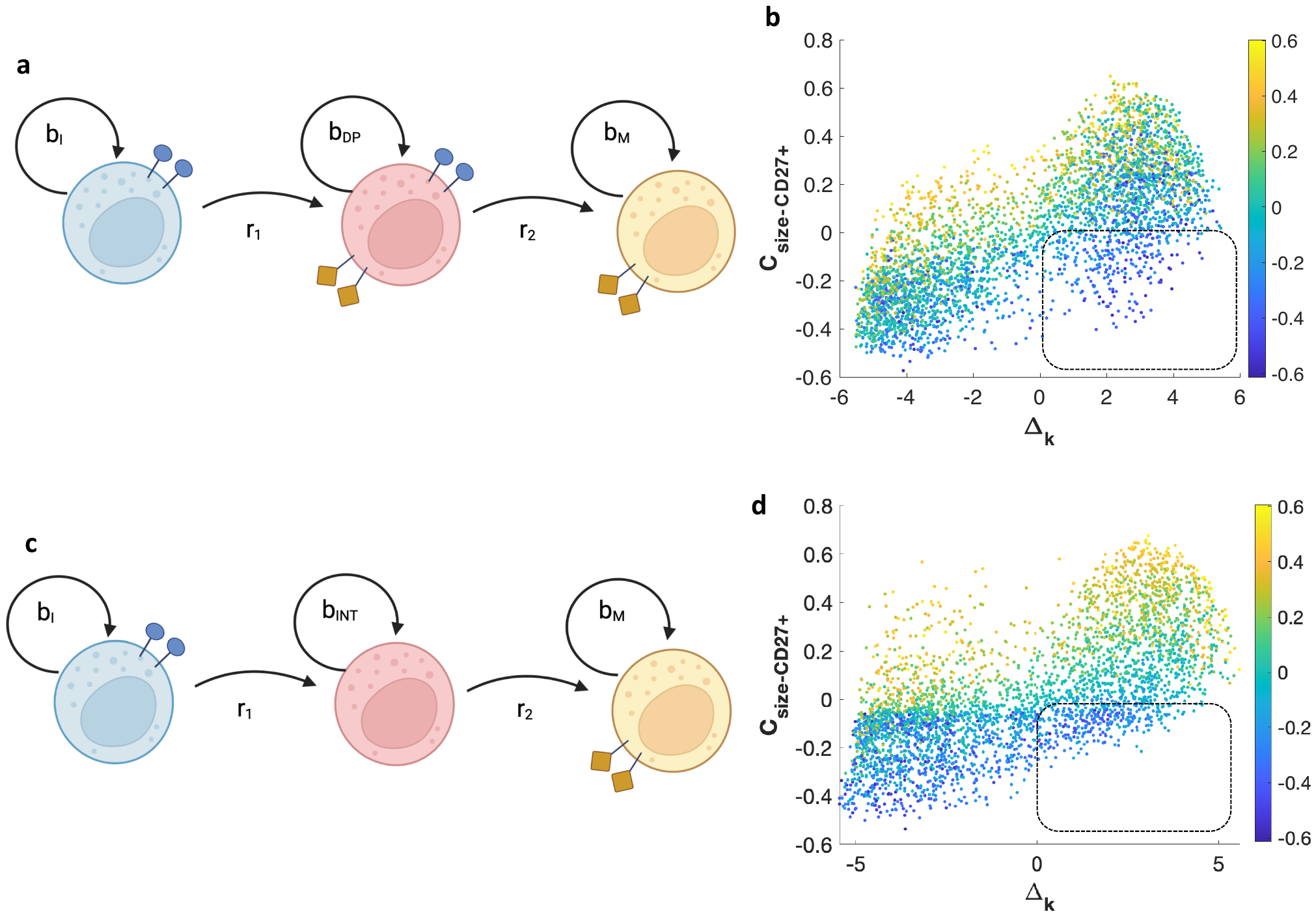
**Figure S5: Three-stage models without death are able to capture negative correlations under the constraint Δ_k_ > 0. (a)** Schematic representation of a three-stage model where the first two stages are CD27+ and the last stage is CD27-. **(b)** A parameter scan for the model shown in (a). Color represents the value *b_I_-b_DP_*. We randomly initialized these simulations with an immature or double-positive cell according to the relative abundances of CD27+CD11b- and CD27+CD11b+ NK cells in the mass cytometry data. Thus, some clones are initialized with an immature cell, and others are initialized with an intermediate double-positive cell. **(c)** Schematic representation of a three-stage model where only the first stage is CD27+ and the last two stages are CD27-. **(d)** A parameter scan for the model shown in (c). Color represents the value *b_I_-b_INT_*.

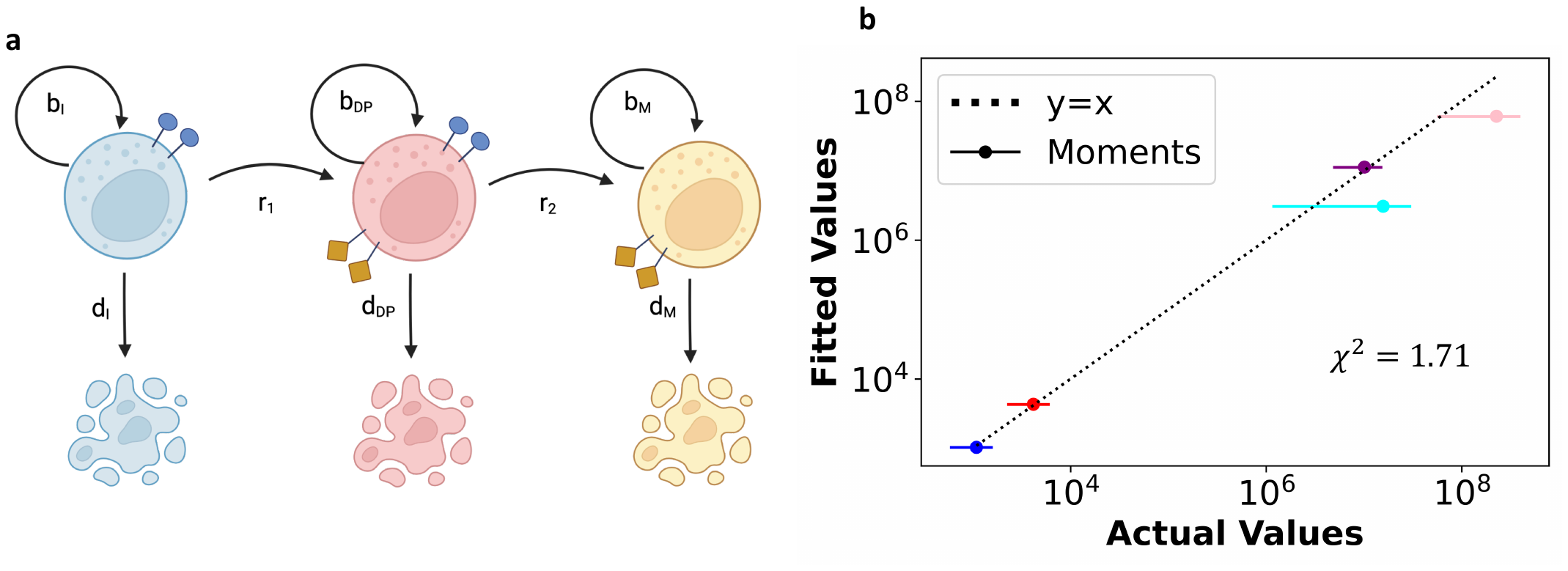

**Figure S6: A three-stage model with two CD27+ stages quantitatively captures clonal burst moments.** **(a)** Schematic representation of a three-stage model where the first two stages are CD27+ and the last stage is CD27-. Each subset undergoes both proliferation and cell death. **(b)** Best fit to the clonal burst data from the model in (a) given the constraints C_size-CD27+_ < -0.2 and Δ_k_ > 0. We used an initial condition for moment solutions and Δ_k_ according to the relative abundances of CD27+CD11b- and CD27+CD11b+ NK cells in the mass cytometry data. Thus, some clones are initialized with an immature cell, and others are initialized with an intermediate double-positive cell.

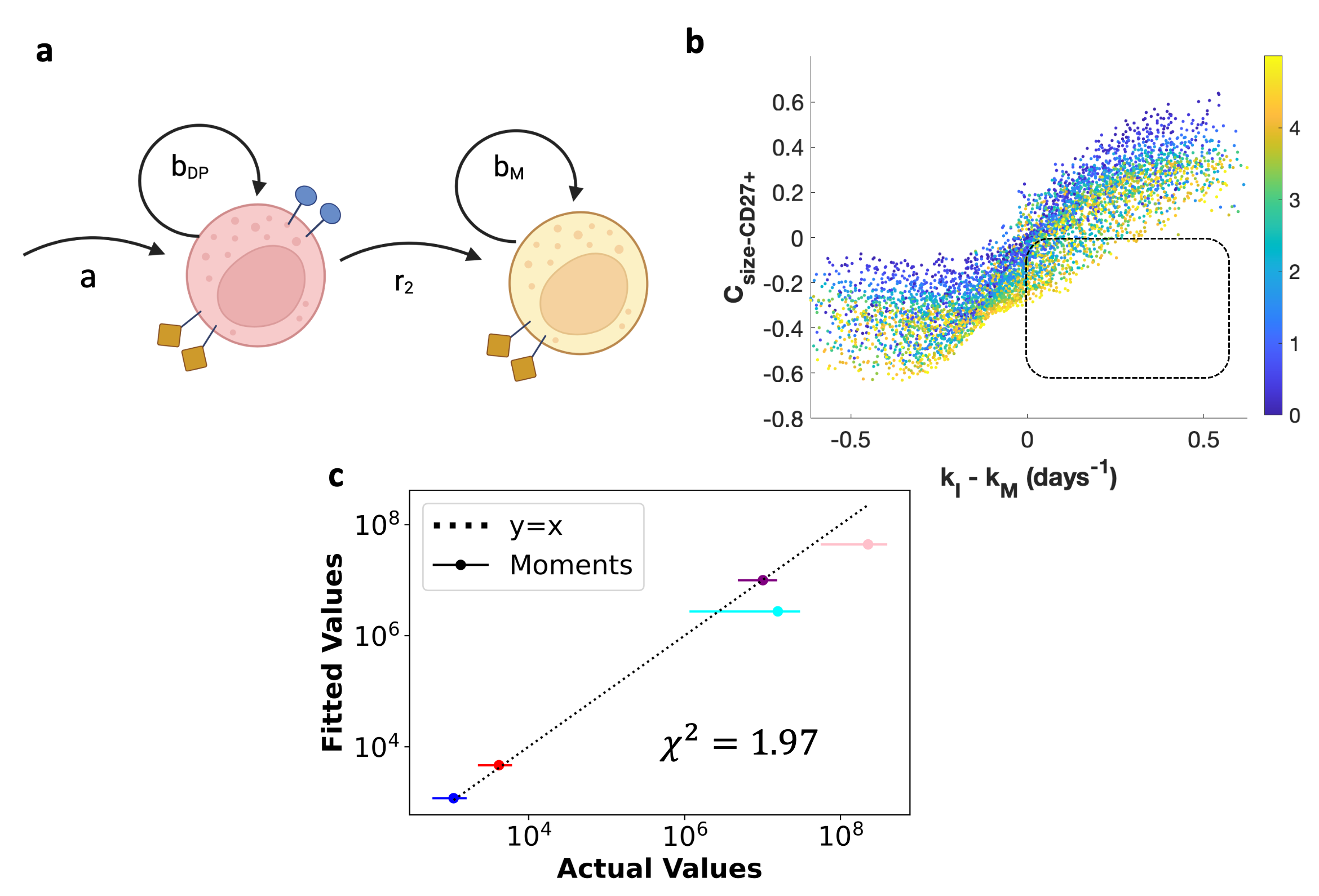

**Figure S7: Mechanistic explanation for how a three-stage model with two CD27+ stages can create negative correlations under constraint Δ_k_ > 0. (a)** Some parameter combinations from the parameter scan in Fig. S5b have *b_I_≅r_1_*, meaning that the change in the first subset is net zero on average. Thus, we investigated a model where this population is held constant to see if negative correlations are still possible. This model is represented here, with a constant inflow *a* into a CD27+ phenotype. This model can be interpreted as having a “stem cell-like progenitor”. **(b)** A parameter scan for the model shown in (a). Color represents the value of *a*. **(c)** Best fit to the clonal burst data from the model in (a) given the constraints C_size-CD27+_ < -0.2 and Δ_k_ > 0.

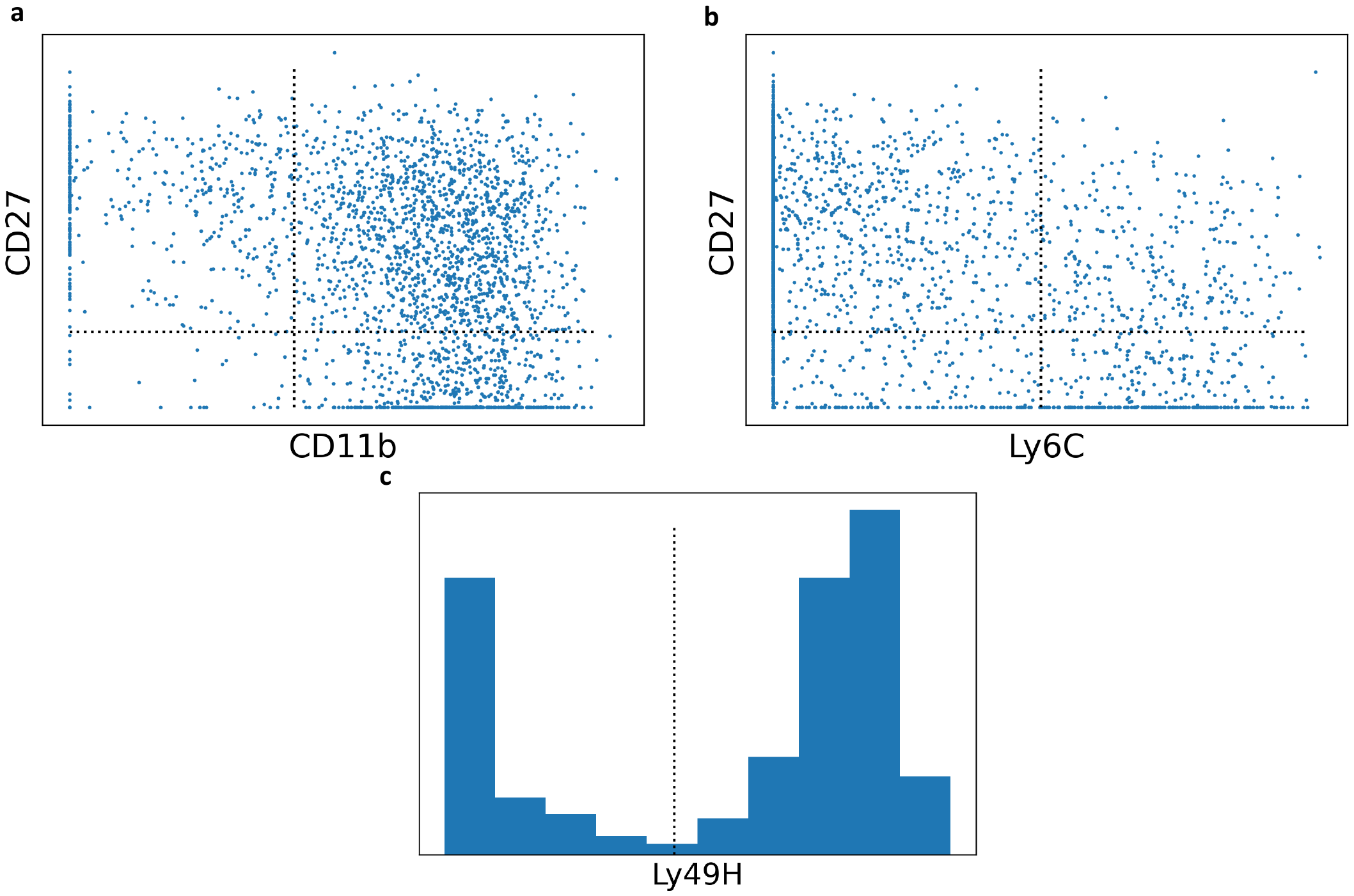

**Figure S8: Gating strategies for mass cytometry data.** All gating strategies are representative from a single mouse’s NK cells. **(a)** Gating strategy for CD27 and CD11b from Ly49H+ NK cells. (**b)** Gating strategy for CD27 and Ly6C from Ly49H+ NK cells. **(c)** Gating strategy for Ly49H.

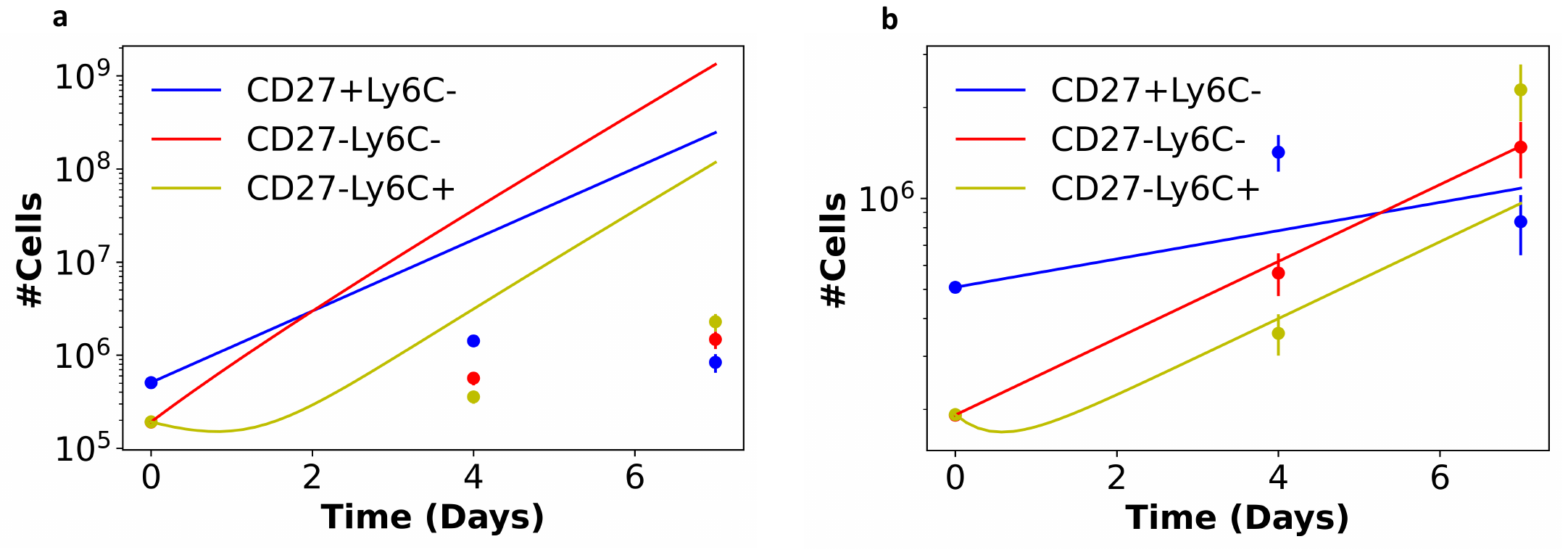

**Figure S9: Supplemental fits to mass cytometry data.** **(a)** Simulated trajectories given the best-fit parameters for the clonal burst data in Table 1. **(b)** Best fit to mass cytometry with constraints that each parameter should be less than 2 and greater than -2 (or greater than 0 for differentiation rates *r_1_* and *r_2_*).

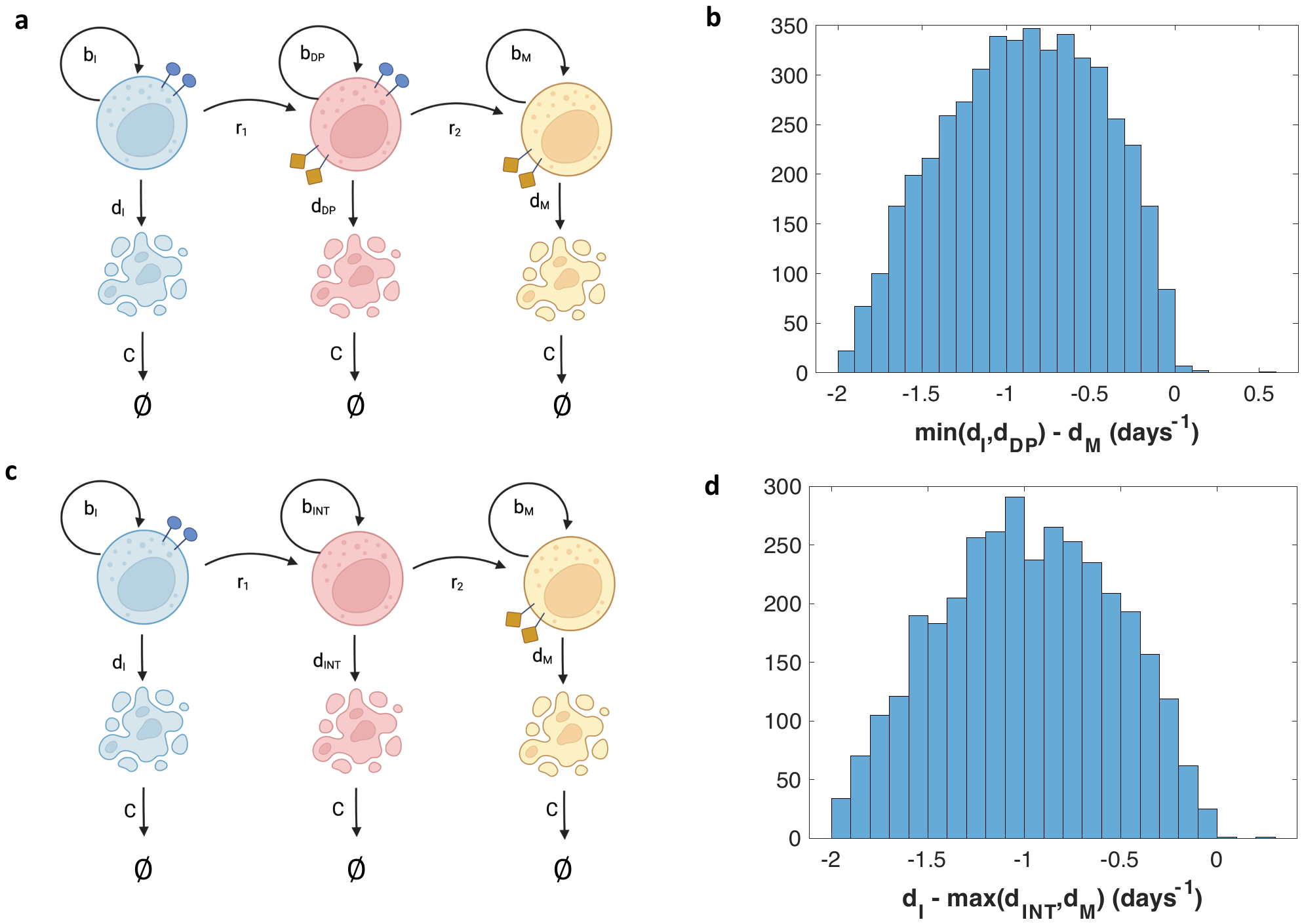

**Figure S10: Models to measure percentage of live cells for each subset demonstrate that percentage of live cells is a good proxy for death rates. (a)** Schematic representation of a three-stage model with two CD27+ subsets, where all dead cells are cleared at a rate *C*. Dead cell abundances are measured, and %live cells can be calculated for CD27+ and CD27- cells by dividing the abundance of live cells by the sum of live and dead cells for each subset. **(b)** A parameter scan given the initial percentages of live cells observed in the experiment in Fig. 3b shows that many parameter configurations generate a higher percentage of living CD27+ cells than that of CD27- cells. Of those parameter configurations, the difference in death rates is shown in this histogram. Because all CD27- death rates are higher, this indicates that a higher %live cells for CD27+ cells requires a higher death rate of CD27- cells. **(c)** Schematic representation of a three-stage model with two CD27- subsets, where all dead cells are cleared at a rate *C*. (**d)** A parameter scan given the initial percentages of live cells observed in the experiment in Fig. 3b shows that many parameter configurations generate a higher percentage of living CD27+ cells than that of CD27- cells. Of those parameter configurations, the difference in death rates is shown in this histogram. Because all CD27- death rates are higher, this indicates that a higher %live cells for CD27+ cells requires a higher death rate of CD27- cells.

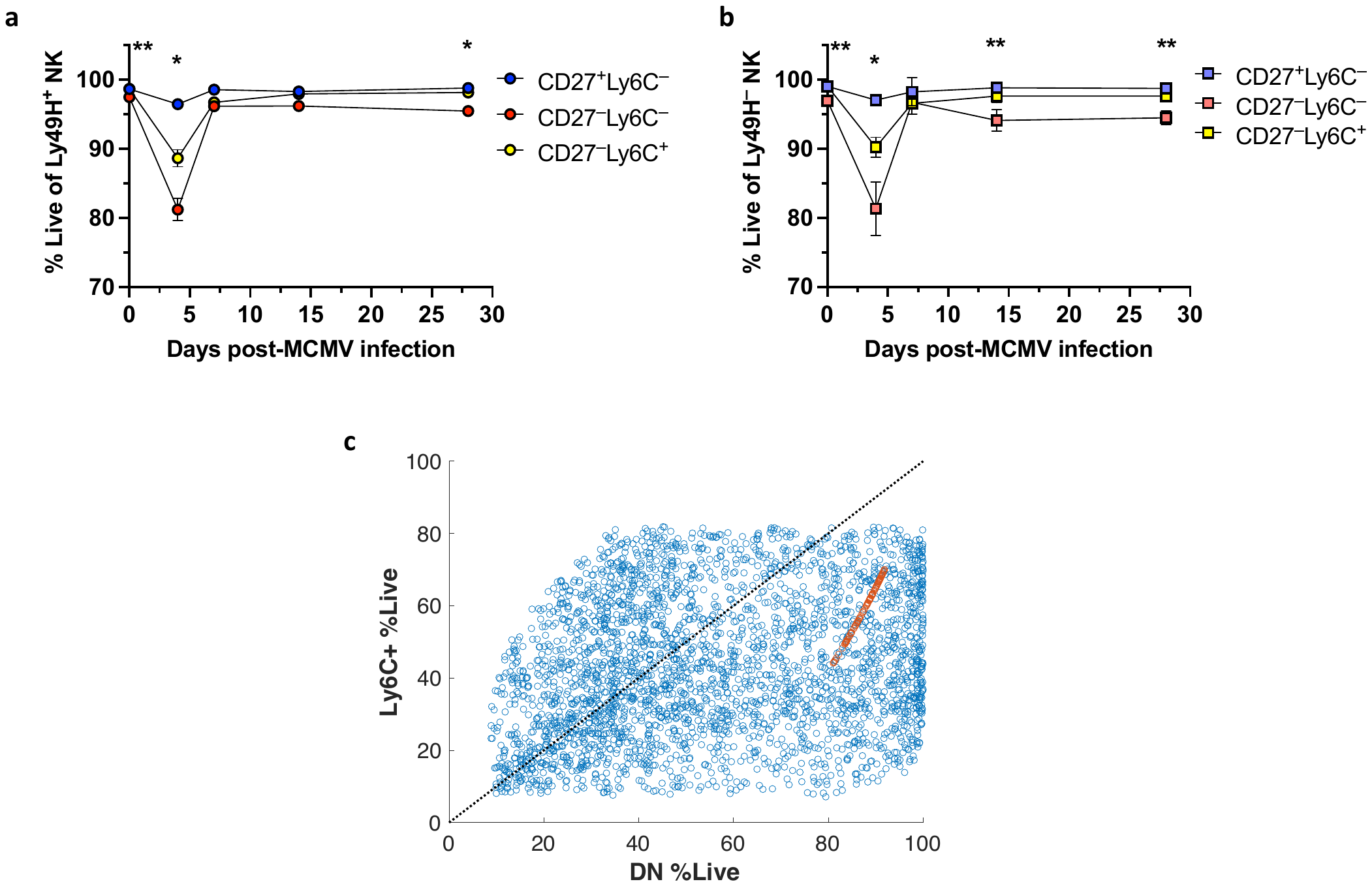

**Figure S11: Both Ly49H+ and Ly49H- CD27- cells show poor survival in *ex vivo* cell culture during the expansion phase response to MCMV infection. (a)** Relative abundances of dead cells for Ly49H+ CD27+Ly6C- (blue), CD27-Ly6C- (red), and CD27-Ly6C+ (yellow) *ex vivo* NK cell subsets as measured by flow cytometry experiment. **(b)** Relative abundances of dead cells for Ly49H- CD27+Ly6C- (blue), CD27-Ly6C- (red), and CD27-Ly6C+ (yellow) *ex vivo* NK cell subsets as measured by flow cytometry experiment. Graphs in (a) and (b) show means ± SEM; *P < 0.033, **P < 0.002 and ***P < 0.001 represent statistically significant difference between CD27^+^ and CD27^–^ NK cells as determined by 2-way ANOVA. **(c)** Parameter scans for determining percentage of live cells for CD27-Ly6C- (DN) and Ly6C+ subsets using the model shown in Fig. S10c. Blue points represent a parameter scan where birth and death rates were varied by random sampling such that the growth rates k_I_ = b_I_-d_I_, k_INT_ = b_INT_-d_INT_, and k_M_ = b_M_-d_M_ are equivalent to the kinetic estimates for endogenous NK cells shown in Table 3. Red points represent randomly sampled dead cell clearance rates with other rates set to those describing adoptive transfer kinetics in Table 1. Black dotted line is *y=x*, and points below this line show a higher percentage of live cells for DN cells than for Ly6C+ cells. Although the estimates in Table 1 don’t accurately capture the higher percentage of live Ly6C+ cells, many configurations of the estimates in Table 3 do.
